# A conformational fingerprint for amyloidogenic light chains

**DOI:** 10.1101/2024.07.12.603200

**Authors:** Cristina Paissoni, Sarita Puri, Luca Broggini, Manoj K. Sriramoju, Martina Maritan, Rosaria Russo, Valentina Speranzini, Federico Ballabio, Mario Nuvolone, Giampaolo Merlini, Giovanni Palladini, Shang-Te Danny Hsu, Stefano Ricagno, Carlo Camilloni

## Abstract

Immunoglobulin light chain amyloidosis (AL) and multiple myeloma (MM) both share the overproduction of a clonal light chain (LC). However, while LCs in MM remain soluble in circulation, AL LCs misfold into toxic soluble species and amyloid fibrils that accumulate in organs, leading to distinct clinical manifestations. The significant sequence variability of LCs has hindered understanding of the mechanisms driving LC aggregation. Nevertheless, emerging biochemical properties, including dimer stability, conformational dynamics, and proteolysis susceptibility, distinguish AL LCs from those in MM under native conditions. This study aimed to identify a conformational fingerprint distinguishing AL from MM LCs. Using small-angle X-ray scattering (SAXS) under native conditions, we analyzed four AL and two MM LCs. We observed that AL LCs exhibited a slightly larger radius of gyration and greater deviations from X-ray crystallography-determined or predicted structures, reflecting enhanced conformational dynamics. SAXS data, integrated with molecular dynamics (MD) simulations, revealed a conformational ensemble where LCs adopt multiple states, with variable and constant domains either bent or straight. AL LCs displayed a distinct, low-populated, straight conformation (termed H state), which maximized solvent accessibility at the interface between constant and variable domains. Hydrogen-deuterium exchange mass spectrometry (HDX-MS) experimentally validated this H state. These findings reconcile diverse experimental observations and provide a precise structural target for future drug design efforts.

## Introduction

Immunoglobulin light-chain (AL) amyloidosis is a systemic disease associated with the overproduction and subsequent amyloid aggregation of patient-specific light chains (LCs) (1–4). Such aggregation may take place in one or several organs, the heart and kidneys being the most affected ones (1). AL originates from an abnormal proliferation of a plasma cell clone that results in LCs overexpression and over-secretion in the bloodstream (1). LCs belonging both to lambda (λ) and kappa (κ) isotypes are associated with AL; however, λ-LCs are greatly overrepresented in the repertoire of AL patients. Specifically, AL-causing LCs (AL-LCs) most often belong to a specific subset of lambda germlines such as *IGLV6* (λ6), *IGLV1* (λ1), and *IGLV3* (λ3) (5–8).

λ-LCs are dimeric in solution with each subunit characterized by two immunoglobulin domains, a constant domain (CL) with a highly conserved sequence and a variable domain (VL) whose extreme sequence variability is the result of genomic recombination and somatic mutations (9–12). VL domains are generally indicated as the key responsible for LC amyloidogenic behavior. The observation that the fibrillar core in most of the structures of *ex-vivo* AL amyloid fibrils consist of VL residues further strengthens this hypothesis (13–17). However, in a recent cryo-electron microscopy (Cryo-EM) structure, a stretch of residues belonging to the CL domain is also part of the fibrillar core, and mass spectrometry (MS) analysis of several *ex vivo* fibrils from different patients indicates that amyloids are composed of several LC proteoforms including full-length LCs (18–21).

Interestingly the overproduction of a light chain is a necessary but not sufficient condition for the onset of AL. Indeed, the uncontrolled production of a clonal LC is often associated with Multiple Myeloma (MM), a blood cancer, but only a subset of MM patients develops AL, thus indicating that specific sequence/biophysical properties determine LC amyloidogenicity and AL onset (12,22–24). To date, the extreme sequence variability of AL-LCs has prevented the identification of sequence patterns predictive of LC amyloidogenicity, however, it has been reproducibly reported that several biophysical properties correlate with LC aggregation propensity. AL-LCs display a lower thermodynamic and kinetic fold stability compared to non-amyloidogenic LCs found overexpressed in MM patients (named hereafter MM-LCs) (12,20–24).

Previous work on LCs has indicated how differences in conformational dynamics can play a role in the aggregation properties of AL-LCs (22–26). Oberti *et al.* have compared multiple λ-LCs obtained from either AL patients or MM patients identifying the susceptibility to proteolysis as the best biophysical parameter distinguishing the two sets (12). Weber *et al.* have shown, using a murine-derived κ-LC, how a modification in the linker region can lead to a greater conformational dynamic, an increased susceptibility to proteolysis, as well as an increased *in vitro* aggregation propensity (25). Additionally, AL-LC flexibility and conformational freedom have also been correlated to the proteotoxicity observed in patients affected by cardiac AL and experimentally verified in human cardiac cells and a *C. elegans* model (27,28). It is noteworthy that the amyloid LCs analyzed in this study were originally purified from patients with cardiac AL amyloidosis.

Here building on this previous work as well on our previous experience on β2-microglobulin, another natively folded amyloidogenic protein (29–33) we aimed at revealing the structural and dynamic determinants of LC amyloidogenicity. With this goal we investigated the native solution state dynamics of multiple λ-LCs by combining MD simulations, SAXS, and HDX-MS. Interestingly, we found a unique conformational fingerprint of amyloidogenic LCs corresponding to a low-populated state (defined as H-state in the following) characterized by extended linkers, with an accessible VL-CL interface and possible structural rearrangements in the CL-CL interface.

## Results

### SAXS suggests differences in the conformational dynamics of amyloidogenic and non-amyloidogenic LC

SAXS data were acquired either in bulk or in-line with size-exclusion chromatography (SEC) for a set of six LCs previously described (cf. **Table 1** and **Methods**). Four of these LCs (referred as H3, H7, H18 in ref. (12) and AL55 in ref. (16)) were identified in multiple AL patients (AL-LC), white two (referred as M7, and M10 in ref. (12)), were identified in MM patients (MM-LC). These LCs cover multiple germlines, with H18 and M7 belonging to the same germline (cf. **Table 1**). The sequence identity is the largest for H18 and M7 (91.6%) while is the lowest for AL55 and M7 (75.2%). A table showing the statistics for all pairwise sequence alignments is reported in **Table S1** in the Supporting Information. For H3, H7, and M7 a crystal structure was previously determined (12) while for H18, AL55, and M10 we obtained a model using either homology modeling (H18 and AL55) or AlphaFold2 (M10). Qualitatively, the SAXS curves in **Figure 1** did not reveal any macroscopic deviation of the solution behavior with respect to the crystal or model conformation. For each LC, we compared the experimental and theoretical curves calculated from the LC structures (cf. **Table 1**) analyzing the residuals and the associated χ^2^. The analysis indicated a discrepancy between the model conformation and the data in the case of the AL-LCs, which was instead not observed in the case of the MM-LCs. For AL-LC, residuals deviate from normality in the low *q* region, suggesting some variability in the global size of the system. Additionally, a weak trend distinguishing AL-LC from MM-LCs could be identified in the radius of gyration (Rg) as derived from the Guinier approximation (cf. **Table 1**). H3, H7, H18, and AL55 have an Rg that is 0.5 to 0.8 Å larger than M7 and M10, a small but statistically significant difference (p-value < 10^−5^). Overall, the SAXS measurements point to less compact and more structurally heterogeneous AL-LCs compared to more compact and structurally homogeneous MM-LCs.

**Table 1.**
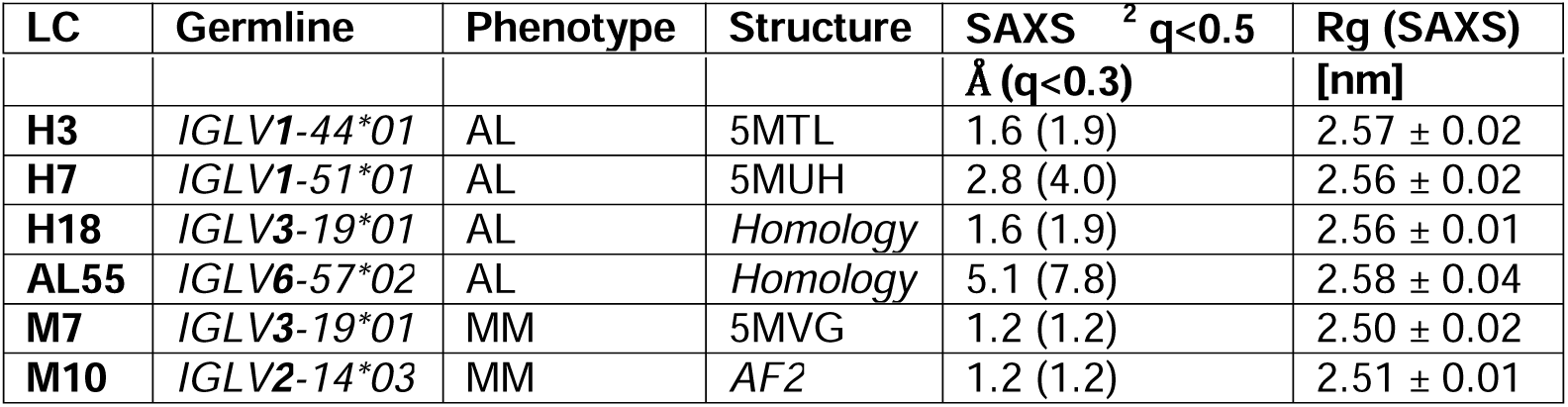
LC systems studied in this work. The table includes information about the germline, phenotype, method to obtain structure, the SAXS curves, and the radius of gyration derived from the SAXS data for all the model proteins studied in this work.

**Figure 1.**
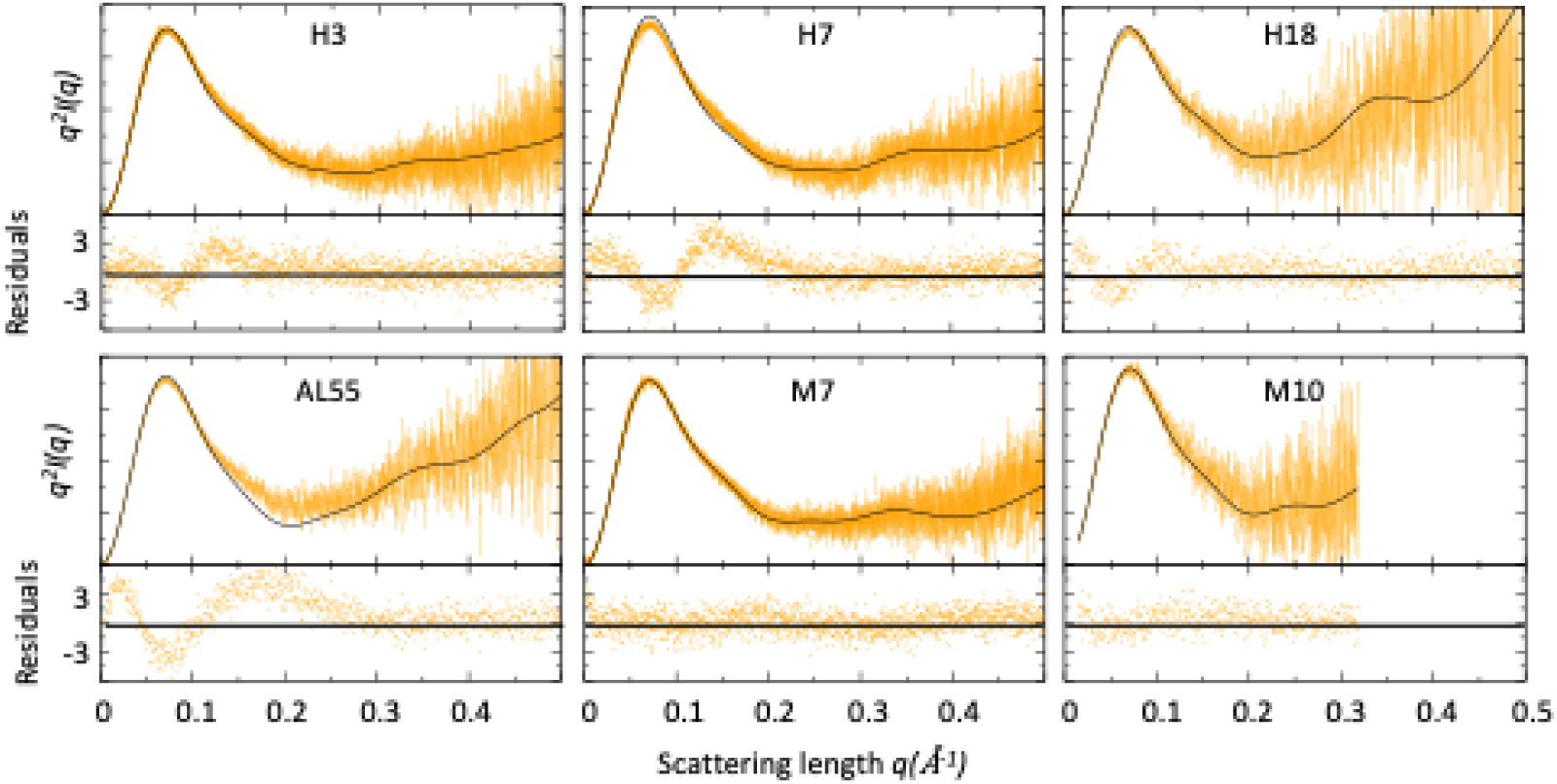
Comparison of SAXS data and single light chain structures. Kratky plots of for AL- and MM light chains. The experimental (orange) and theoretical (black) curves (top panels) and the associated residuals (bottom panels) indicate that AL-LC solution behavior deviates from reference structures more than MM-LC. SAXS was measured as follow: H3 measured in bulk (Hamburg), 3.4 mg/ml; H7 measured in bulk (Hamburg), 3.4 mg/ml; H18 measured by SEC-SAXS (ESRF) with the injection concentration of at 2.8 mg/ml; AL55 measured in bulk (ESRF), 2.6 mg/ml; M7 measured in bulk (Hamburg), 3.6 mg/ml; M10 measured by SEC-SAXS with the injection concentration of 6.7 mg/ml (ESRF). Theoretical SAXS curves were calculated using crysol (54). Log-log plots are shown in Fig. S1 in the Supporting Information.

### MD simulations reveal a conformational fingerprint for amyloidogenic light chains

To investigate the conformational dynamics of the six LCs we performed Metadynamics Metainference (M&M) MD simulations employing the SAXS curves (*q<0.3* Å) as restraints (cf. **Material and Methods**) (34–37). Metainference is a Bayesian framework that allows the integration of experimental knowledge on-the-fly in MD simulations improving the latter while accounting for the uncertainty in the data and their interpretation. Metadynamics is an enhanced sampling technique able to speed up the sampling of the conformational space of complex systems. The combination of SAXS and MD simulations has been shown to be effective for multi-domain proteins as well as for intrinsically disordered proteins (38–40).

For each LC, we performed two independent M&M simulations coupled by the SAXS restraint, accumulating around 120-180 µs of MD per protein (cf. **Material and Methods** and **Table 2**). The resulting conformational ensembles resulted in a generally improved agreement with the SAXS data employed as restraints (cf. **Table 2** and **Figure 2A**). To investigate differences in the LC local flexibility we first analyzed the root mean square fluctuations (RMSF) for the CL and VL separately, averaging over the chains and the replicates, **Figure 2B**. The RMSF indicates comparable flexibility in most of the regions, with differences localized in the termini and in some loops. The VL of amyloidogenic LCs are generally more flexible than the MM ones, but this may be associated with the lengths of their complementarity-determining regions (CDRs). Indeed, M10 has the longest and most flexible CDR1 and the shortest and least flexible CDR3. Unexpectedly, there are some differences also in the CL domains. Here, in **Figure 2B**, the AL-LCs are always more flexible in at least one region even if these differences are relatively small. Overall, the RMSF does not provide a clear indication to differentiate AL and MM-LCs. To provide a global description of the dynamics of the six LC systems, we then introduced two collective variables, namely the elbow angle, describing the relative orientation of VL and CL dimers, and the distance between the VL and CL dimers center of mass, illustrated in **Figure 2C**.

**Table 2.** Metainference simulations performed in this work for the 6 systems. For each metainference simulation is reported the simulation time per replica with the number of replicas; the chi-square of the resulting conformational ensemble with the experimental SAXS curve, the range q<0.3 Å is the one used as restraint in the simulation; the average radius of gyration with error estimated by block averaging.

| LC code | Simulation | Length per replica<br>[ns] (# replicas) | SAXS $\chi^2$ $q<0.5$<br>$\text{\AA} (q<0.3)$ | Average<br>Rg [nm] |
| --- | --- | --- | --- | --- |
| **H3** | M&M 1 | 1,530 (60) | 1.2 (1.1) | $2.56 \pm 0.02$ |
| | M&M 2 | 1,520 (60) | 1.2 (1.1) | $2.56 \pm 0.02$ |
| **H7** | M&M 1 | 1,627 (60) | 1.1 (1.2) | $2.54 \pm 0.04$ |
| | M&M 2 | 1,545 (60) | 1.1 (1.2) | $2.54 \pm 0.05$ |
| **H18** | M&M 1 | 1,643 (60) | 1.4 (1.6) | $2.53 \pm 0.03$ |
| | M&M 2 | 1,529 (60) | 1.4 (1.7) | $2.54 \pm 0.03$ |
| **AL55** | M&M 1 | 1,545 (60) | 2.7 (3.0) | $2.59 \pm 0.02$ |
| | M&M 2 | 1,591 (60) | 2.5 (2.7) | $2.58 \pm 0.05$ |
| **M7** | M&M 1 | 1,623 (60) | 1.2 (1.2) | $2.52 \pm 0.03$ |
| | M&M 2 | 1,530 (60) | 1.1 (1.2) | $2.52 \pm 0.03$ |
| **M10** | M&M 1 | 987 (60) | 1.1 (1.1) | $2.53 \pm 0.07$ |
| | M&M 2 | 995 (60) | 1.1 (1.1) | $2.54 \pm 0.05$ |

**Figure 2.**
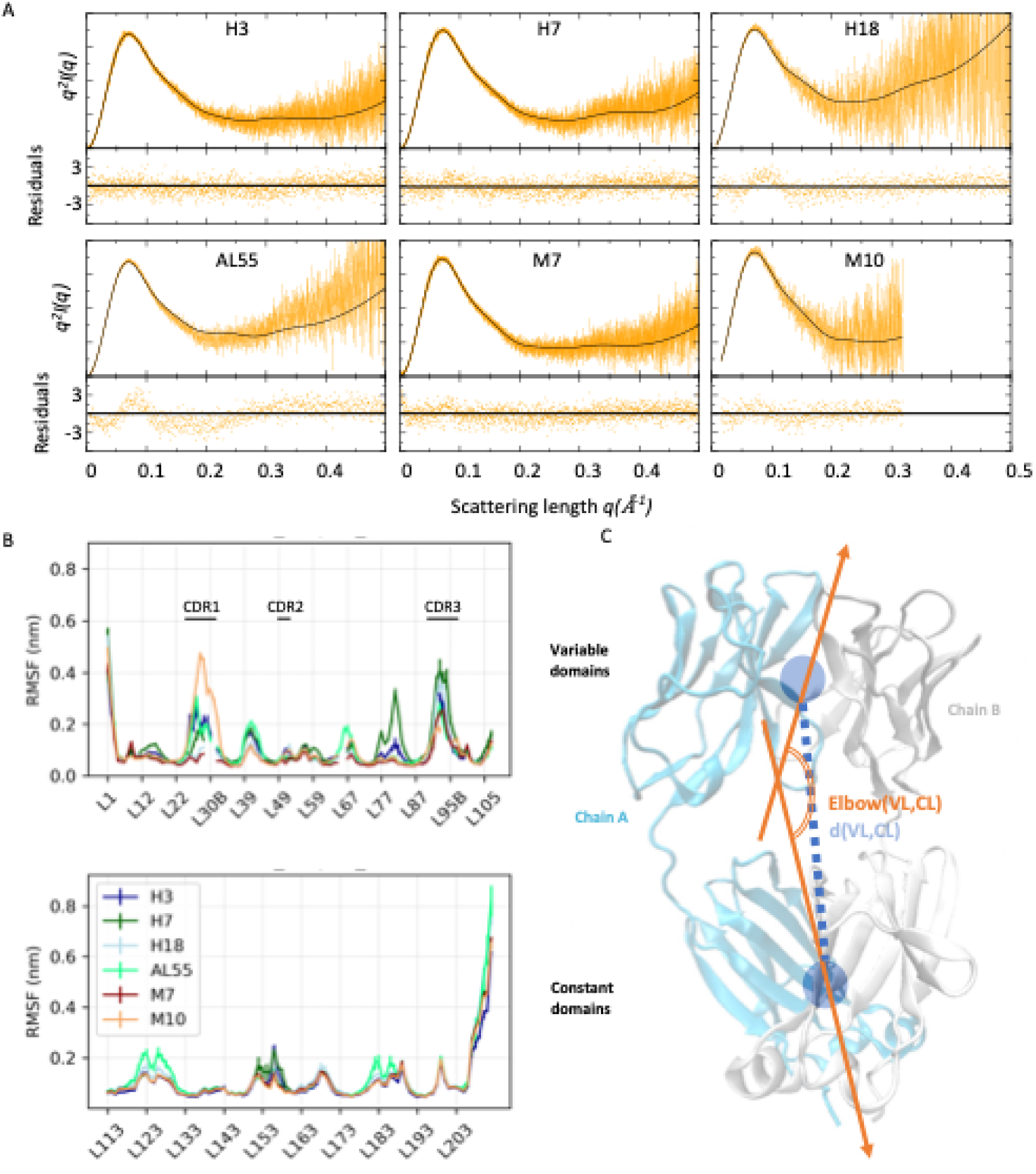
Light chain SAXS driven MD simulations. (A) Kratky plots and associated residuals (bottom panels) comparing experimental (orange), and theoretical (black) curves obtained by averaging over the metainference ensemble for H3, H7, H18, AL55, M7 and M10, respectively. Theoretical SAXS curves were calculated using crysol (54). (B) Residue-wise root mean square fluctuations (RMSF) obtained by averaging the two Metainference replicates and the two equivalent domains for the six systems studied. The top panel shows data for the variable domains, while the bottom panel shows data for the constant domain. Residues are reported using *Chothia and Lesk* numbering (51). (C) Schematic representation of two global collective variables used to compare the conformational dynamics of the different systems, namely the distance between the center of mass of the VL and CL dimers and the angle describing the bending of the two domain dimers.

In **Figure 3**, we report the free energy surfaces (FES) obtained from the processing of the two replicates of each LC as a function of the elbow angle and the CL-VL distance calculated from their center of mass. The visual inspection of the FES indicates converged simulation: in all cases, the replicates explore a comparable free-energy landscape with comparable features. All six LC FES share common features: a relatively continuous low free energy region along the diagonal, spanning configurations where the CL and VL are bent and close to each other (state L_B_), and configurations where the CL and VL domains are straight and at relative distance between 3.4 and 4.1 nm (state L_S_). A subset of LCs, namely H18, M7, and AL55, display conformations where the domains are straight in line (elbow angle greater than 2.5 rad) and in close vicinity, with a relative distance between the center of mass of less than 3.4 nm (state G). Of note, H18 and M7 belong to the same germline, letting us speculate that this state G may be germline specific. Most importantly, only the AL-LCs display configurations with CL and VL straight in line but well separated at relative distances greater than 4.1 nm, this state H seems to be a fingerprint specific for AL-LCs. A set of configurations exemplifying the four states is reported in **Figure 3**. The estimates of the populations for the four states L_B_, L_S_, G, and H are reported in **Table 3**. The quantitative analysis indicates that, within the statistical significance of the simulations, states L_B_ and L_S_ represent in all cases most of the conformational space. In the case of H18, AL55, and M7, the compact state G is also significantly populated (10-34%). The state H, associated with amyloidogenic LCs, is populated between 5 and 10% in H3, H7, H18, and AL55 and less than 1% in M7 and M10.

**Figure 3.**
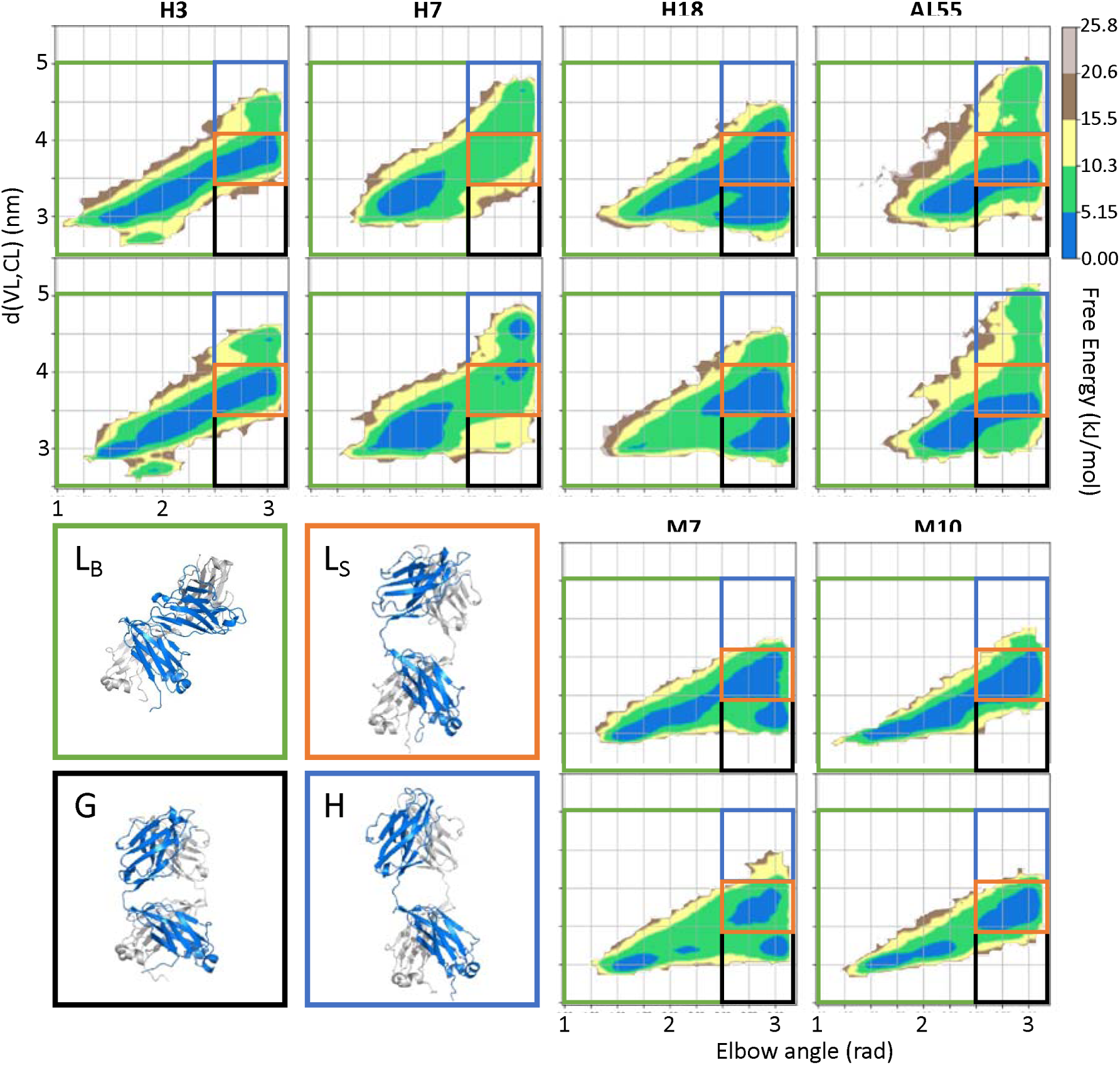
Free Energy Surfaces (FESes) for the six light chain systems under study by Metadynamics Metainference MD simulations. For each system, the simulations are performed in duplicate. The x-axis represents the elbow angle indicating the relative bending of the constant and variable domains (in radians), while the y-axis represents the distance in nm between the center of mass of the CL and VL dimers. The free energy is shown with color and isolines every 2k_B_T corresponding to 5.16 kJ/mol. On each FES are represented four regions (green L_B_-state, red L_S_-state, blue H-state, and black G-state) highlighting the main conformational states. For each region, a representative structure is reported in a rectangle of the same color.

**Table 3.** Populations of the four states shown in Figure 3 resulting from the two independent Metadynamics Metainference simulations performed for each of the 6 LCs. The population of the H state, which we supposed to be a fingerprint specific for AL-LCs, is in bold.

| % | **H3** | **H7** | **H18** | **AL55** | **M7** | **M10** |
| --- | --- | --- | --- | --- | --- | --- |
| **L<sub>B</sub>** | 62.0±0.4 | 72.3±2.5 | 22.9±2.8 | 48.3±0.1 | 46.8±0.1 | 48.4±0.3 |
| **L<sub>S</sub>** | 33.0±0.2 | 15.2±2.7 | 38.4±2.1 | 32.5±2.0 | 35.0±0.1 | 49.1±1.5 |
| **G** | 0.2±0.1 | 0.8±0.4 | 33.8±3.8 | 10.5±1.4 | 17.6±0.1 | 1.8±0.6 |
| **H** | **4.8±0.5** | **11.7±0.5** | **5.0±1.0** | **8.7±0.5** | 0.6±0.2 | 0.8±0.6 |

To identify additional differences between the conformations observed in state H and the rest of the conformational space, we focused our attention on the VL-VL and CL-CL dimerization interfaces. In **Figure 4**, we showed the free energy as a function of the distance between the CL domains versus the distance between the VL domains for each of the four states for one of the two simulations performed on H3; the same analysis for all other simulations is shown in **Figures S2 to S7** in the Supporting Information. From the comparison of the FESes, it is clear that only in the conformations corresponding to the state H do the CL-CL dimers display an alternative configuration. In the case of H3, the CL-CL domains in the H state are characterized by a shift towards configurations characterized by a larger distance, the same is observed in the case of H18 and AL55, while in the case of H7 the H state is characterized by a smaller distance between the CL domains.

**Figure 4.**
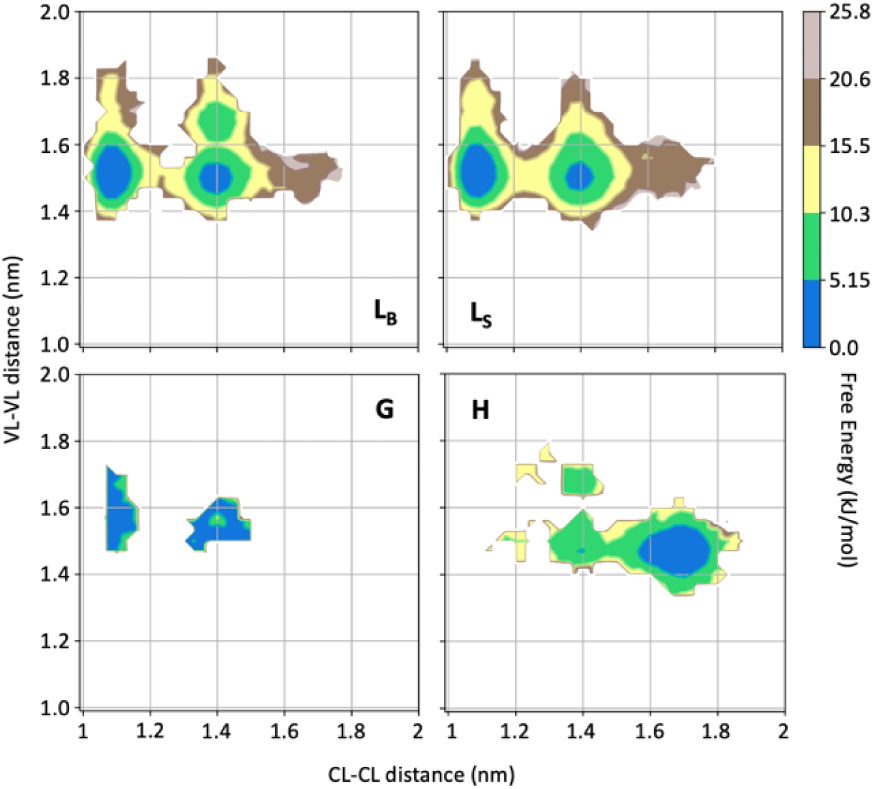
FESes for the four substates identified in Figure 3 in the case of the first H3 Metainference simulation. The x-axis shows the distance between the centers of mass of the constant domains, while the y-axis shows the distance between the centers of mass of the variable domains. The free energy is shown with color and isolines every 2k_B_T corresponding to 5.16 kJ/mol.

Our conformational ensembles allowed us to hypothesize a conformational fingerprint for AL proteins, namely the presence of a weakly but significantly populated state (H) characterized by a more extended quaternary structure, with VL and CL dimers well separated, and with perturbed CL-CL interfaces.

### HDX independently validates the amyloidogenic LC conformational fingerprint

To gain further molecular insight into how the dynamics of the tertiary and quaternary structures can be differentiated in AL- and MM-LCs, HDX-MS was performed on our set of proteins. HDX-MS probes the protein dynamics by monitoring the hydrogen-to-deuterium uptake over time and the obtained data well complement structural, biophysical, and computational data. Four LCs from our set (H3, H7, AL55, and M10) yielded good peptide sequence coverages of 98.6, 92.5, 98.6, and 99.1%, respectively, with redundancy of > 4.0 (**Figures S8A-S11A,** and **Table S2** in the Supporting Information). H18 and M7 were not included in this analysis due to their poor sequence coverage and were not further investigated. Due to sequence heterogeneity among our proteins, the common peptide analyses were not performed. Therefore, the relative deuterium exchange at different HDX time points of 0.5 to 240 min with respect to zero exchange time was used to compare the dynamics of individual proteins. The average uptake at different time points for selected regions including residues 34-50 and 152-180 were included in **Figure S12** in the Supporting Information. The individual peptide mapping at all HDX time points is given in **Figure S13** in the Supporting Information. The deuterium uptakes at 30 min HDX time showed the most pronounced differences between different proteins, which were chosen to illustrate the key structural features in the main figure panel (**Figure 5**).

**Figure 5.**
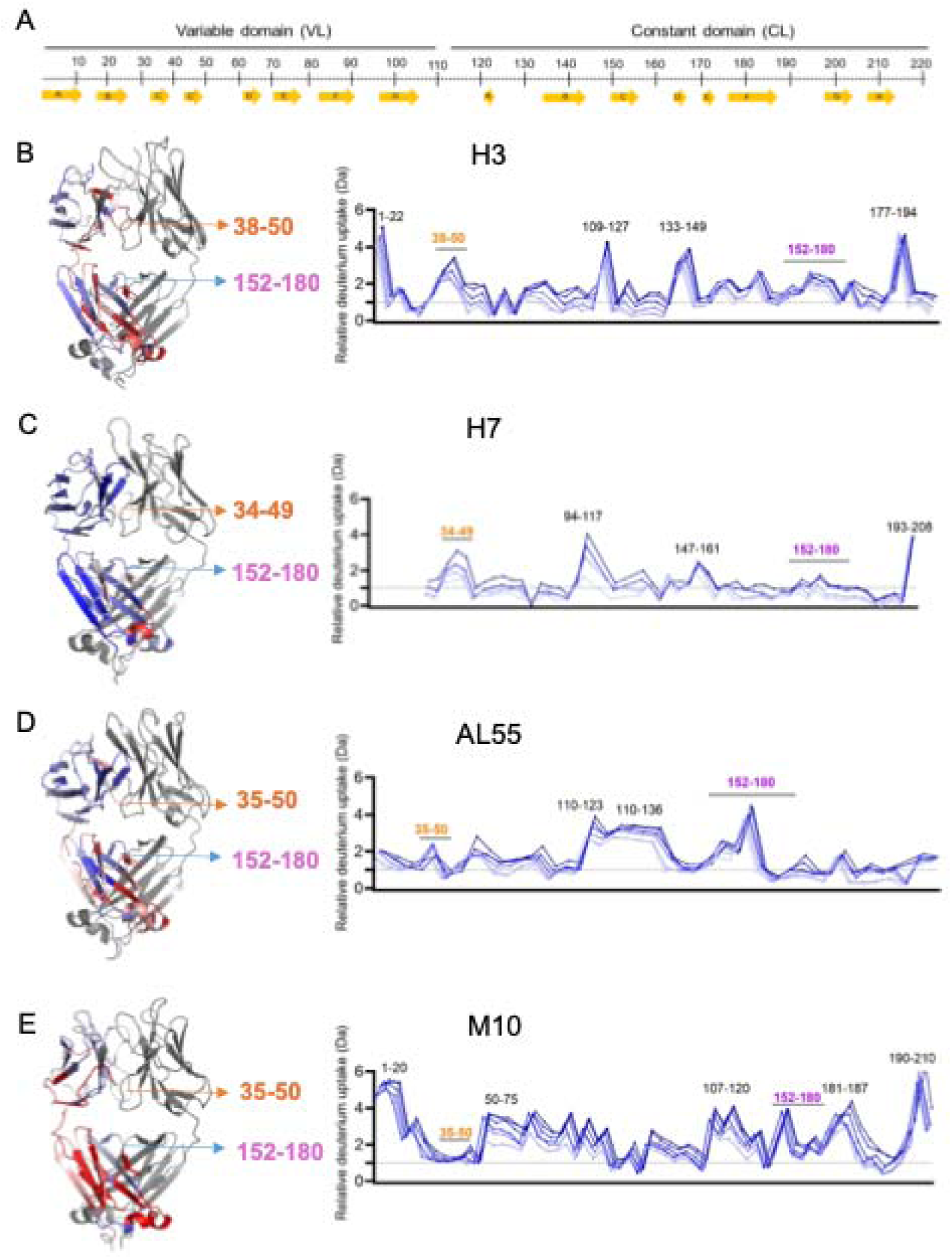
HDX-MS analysis. (A) The top panel represents the simplified presentation of the primary structure of an LC including variable domain (VL) and constant domain (CL). The location of β-strands according to *Chothia and Lesk* (51). (B) The structural mapping and butterfly plot of relative deuterium uptake (Da) of H3. The chain A of H3 structure is colored in a gradient of blue-white-red for an uptake of 0 to 30% at an exchange time of 30 min. The chain B is shown in gray (right hand panel). The butterfly plot showing relative deuterium uptake at all time points from 0.5 to 240 min on a gradient of light to dark blue (left-hand panel). (C, D, E) are the figures corresponding to H7, AL55, and M10, respectively with the same color coding as in figure B. The overall sequence coverage for all proteins was > 90% with a redundancy of > 4.0. The VL-VL domains interface peptides covering amino-acid residues 34-50 are labelled in orange while CL-CL interface region containing 152-180 amino acids are labelled in magenta. Collectively, they form VL-CL interface which is important to define H-state.

HDX-MS analysis revealed subtle structural dynamics of the individual proteins. The most significant difference between the AL and MM-LCs is observed for residues 34-50, which are part of both the VL-VL dimerization interface and, more importantly in the context of this work, the CL-VL interface. These residues show significantly higher deuterium uptakes in all H-proteins, with H3 being the highest, implying that AL-LCs dimeric interfaces (VL-VL and CL-VL) are more dynamic and hence significantly more destabilized than in M10 (**Figure 5, Figure S12** in the Supporting Information). The highly dynamic VL-VL interface of H3 also correlates well with its open VL-VL interface in a crystal structure (PDB 8P89), which houses two nanobodies interacting with each VL in a dimeric structure (28). On the other hand, residues 54-70, which are not part of either interface, show a higher deuterium uptake and hence more dynamics in M10 protein, which may be a result of redistribution of dynamics in the regions away from the rigid VL-VL interface to stabilize the overall structure as also observed for other proteins previously (41,42) (**Figure 5, Figure S12**). In contrast, the VL-CL hinge regions (residues 110-120) show homogenously high flexibility in all the proteins except H7 indicating their higher accessible surface areas (**Figure 5**). As expected, the CL domains from residues 170-200 also show a similar pattern of average deuterium uptakes and hence flexibility in AL- and MM-LCs, with a minor difference contributed by the rigid VL-CL interface containing residues 165-180 (**Figures S8B-S11B, S12** in the Supporting Information). This region shows significantly less deuterium uptake (rigid) in both AL and M proteins when compared to other peptides in the CL domain (**Figures S8B-S11B** in the Supporting Information). However, comparing the average uptake for this region (residue 152-180) between AL and M proteins shows that H3 and AL55 have higher uptake than M10. In contrast, H7 is an exception with the lowest deuterium uptake in this region (**Figure S12 Panel 152-180** in the Supporting Information). These data are particularly interesting in the light of our simulations. The dimeric conformations identified in the H state, **Figure 3**, are characterized by higher accessibility for the CL-VL interface, which is in agreement with the increased accessibility for the region 34-50 on the VL and 159-180 in the CL observed in the HDX-MS analysis. Notably, the H state of H7 is the only one in which the CL-CL interface is remarkably compact (see **Figure S3** in the Supporting Information), consistent with the lower HDX for residues 152-180 observed in H7. Overall, the HDX-MS data provide an independent validation of the H-state predicted from our conformational ensembles

## Discussion

Understanding the molecular determinants of AL amyloidosis has been hampered by its high sequence variability in contrast to its highly conserved three-dimensional structure (8). In this work, building on our previous studies highlighting susceptibility to proteolysis as a property that can discriminate between AL-LC and MM-LC, as well as the role of conformational dynamics in protein aggregation, we characterized LC conformational dynamics under the assumption that AL-LC proteins, despite their sequence diversity, may share a property that emerges at the level of their dynamics. We combined SAXS measurements with MD simulations under the integrative framework of Metainference to generate conformational ensembles representing the native state conformational dynamics of 4 AL-LC and 2 MM-LC. While SAXS alone already indicated possible differences, its combination with MD allowed us to observe a possible low-populated state, which we refer to as state H, characterized by well-separated VL and CL dimers and a perturbed CL-CL interface, which is significantly populated in AL-LCs while only marginally populated in MM-LCs. Notably, high-energy states associated with amyloidogenic proteins have been previously identified in the case of SH3 (43) and β2m (32). Here, HDX measurements allowed us to independently validate this state by observing increased accessibility in CL-VL interface regions. Importantly, given the limited size of our LC set, we observed in reference (44) that a comparison of lambda-6 LCs by HDX-MS revealed differences between AL and MM LCs in a region overlapping with the one reported here (residues 35–50). Specifically, this region exhibited increased deuterium uptake in AL compared to MM proteins. A T32N substitution was shown to mitigate aggregation propensity and reduce deuterium uptake in the AL protein, while the reverse substitution, N32T, applied to the germline sequence, increased it (44). Finally, the H-state observed here is comparable to that previously observed for an aggregation-prone mutation in a monomeric kappa LC (25).

Having established a conformational fingerprint for AL-LC proteins, it would be tempting to identify possible mutations that could be associated with the presence of the H state. Comparing the sequences and structures of M7 and H18, both of which belong to the *IGLV3-19*01* germline, we can identify a single mutation, A40G, that could easily be associated with the appearance of the H state in H18. This mutation is located in the 37-43 loop, which H/D exchange showed to be more accessible in our three AL-LCs than in our MM-LC (see **Figure 5 and Figure S12**), and it breaks a hydrophobic interaction with the methyl group of T165, as observed in the crystal structure of M7 (PDB 5MVG and **Figure S13**), potentially making T165 more accessible in H18 than in M7 (see **Figure 5, and Figure S12**). Comparing the H18 and M7 sequences with the germline reference sequence, we see that position 40 in *IGLV3-19*01* is a glycine (see **Figure S13**). This would suggest the intriguing interpretation that the G40A mutation in M7 may increase the interdomain stability compared to the germline sequence, making it less susceptible to aggregation. However, it should also be noted that while this framework position is a glycine in H3, H7, and AL55, it is also a glycine in M10. Previous research has often focused on identifying, on a case-by-case basis, the key mutations that may be considered responsible for the emergence of the aggregation propensity, under the assumption that such aggregation propensity should not be present in germline sequences, but this assumption may be misleading given the observation that few germlines are strongly overrepresented in AL, suggesting that these starting germline sequences may be inherently more aggregation-prone than the germline genes that are absent or rarely found in AL patients. More generally, by comparing our AL-LC sequences with their germline references (**Figures S14 to S19** in the Supporting Information), we observe that all mutations fall exclusively in the variable domain, allowing us to exclude for these systems a direct role for residues in the linker region, as observed in ref (25,45), or in the constant domain, as observed in ref (46). Many mutations fall in the CDR regions, as expected, but others are found in the framework regions, both near the dimerization interface and in other regions of the protein. Somatic mutations in the dimerization interfaces have been identified previously as possibly responsible for toxicity of AL-LC (47). Regarding mutations in the CDRs, it has been suggested that AL-LC proteins may exhibit frustrated CDR2 and CDR3 loops, with few key residues populating the left-hand alpha helix or other high-energy conformations (48), resulting in the destabilization of the VL. In **Figures S14 to S19** in the Supporting Information, we have analyzed the Ramachandran plot obtained from our conformational ensembles, focusing only on those residues that most populate the left-hand alpha helix region, which are marked with a red circle symbol and whose Ramachandran is reported. Our data indicate the presence of residues populating left-hand alpha regions in the Ramachandran plot, but these are also found in the case of MM-LCs, so our simulations do not allow to confirm or exclude this mechanism in our set of protein systems.

In conclusion, our study provides a novel, complementary, perspective on the determinants of the misfolding propensity of AL-LCs that we schematize in Figure 6. The identification of a high-energy state, with perturbed CL dimerization interfaces, extended linkers, and accessible regions in both the VL-CL and VL-VL interfaces may be the common feature interplaying with specific properties shown by previous work including the direct or indirect destabilization of both the VL-VL and CL-CL dimerization interfaces (22,23,46–50). Our conformational fingerprint is also consistent with the observation that protein stability does not fully correlate with the tendency to aggregate, whereas susceptibility to proteolysis and conformational dynamics may better to capture the differences between AL-LC and MM-LC. In this context, our data allow us to rationally suggest that targeting the constant domain region at the CL-VL interface, which is more labile in the H state, maybe a novel strategy to search for molecules against LC aggregation in AL amyloidosis.

**Figure 6.**
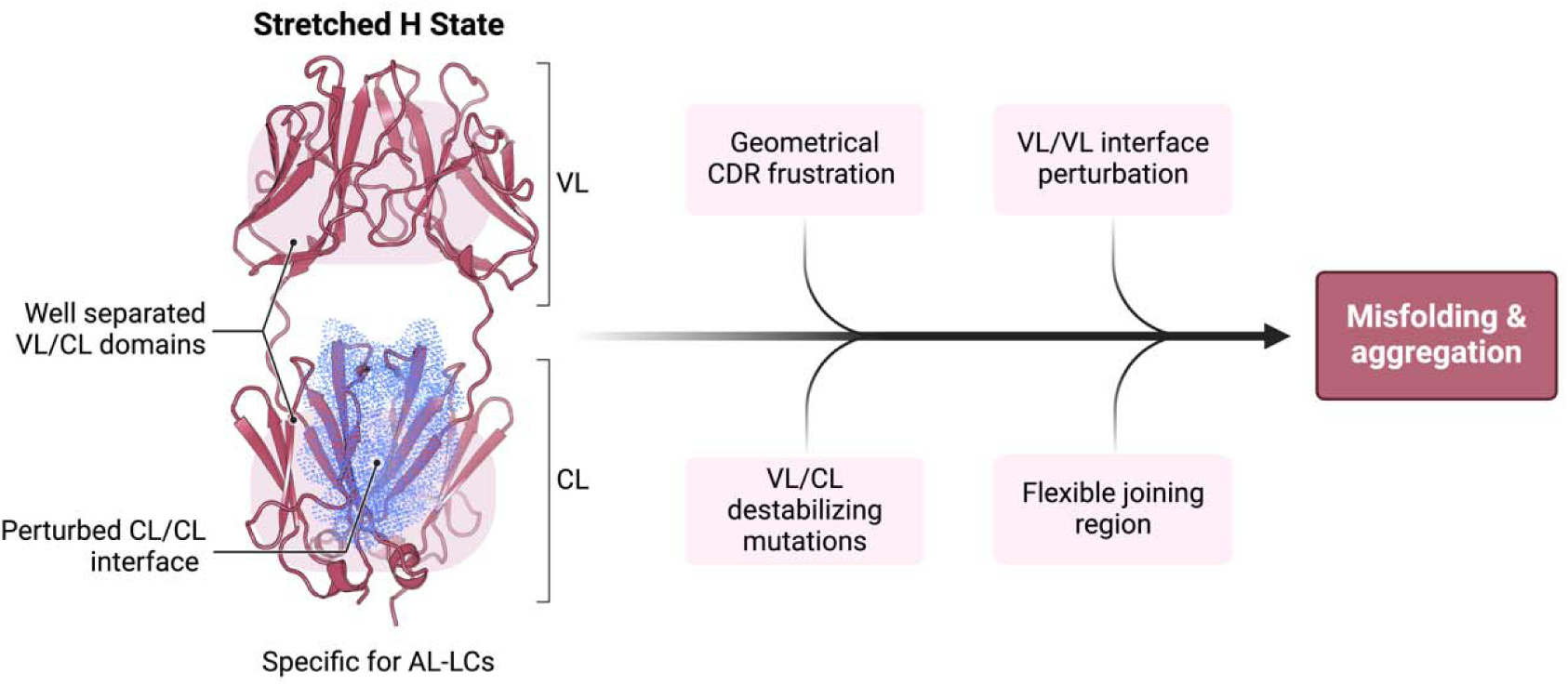
Schematic representation summarizing our findings in the context of previous work on the biophysical properties of amyloidogenic light chains. We propose that the H state is the conformational fingerprint distinguishing AL LCs from other LCs, which together with other features contributes to the amyloidogenicity of AL LCs.

## Materials and Methods

### LC production and purification

Recombinant AL- (H3, H7, H18, and AL5) and M- (M7, M10) proteins were produced and purified from the host *E. coli* strain BL21(DE3). Firstly, the competent BL21(DE3) cells were transformed with plasmid pET21(b+), which contains genes encoding H3, H7, H18, AL55, M7, and M10 proteins. The transformed cells were selected for each plasmid by growing them on LB agar plates containing the antibiotic ampicillin at a final concentration of 100 µg/ml. For over-expression of protein, one colony was picked from each plate and grown overnight in 20 ml of LB broth containing ampicillin at a final concentration of 100 µg/ml. The overnight-grown cells were then used to inoculate a secondary culture in one liter of LB broth. The cells were grown until the turbidity (OD_600nm_) reached between 0.6-0.8 and protein expression was subsequently induced by adding 0.5 mM isopropyl-β-D-thiogalactopyranoside (IPTG) for 4 h. The bacterial cells containing overexpressed LCs were then harvested using a Backman Coulter centrifuge at 6000 rpm for 20 min at 4 °C. All the proteins were overexpressed as inclusion bodies. For protein purification, the inclusion bodies were isolated by cell lysis induced by sonication. The purification of inclusion bodies was performed by washing them with buffer containing 10 mM Tris (pH 8) and 1% triton X 100. The purified inclusion bodies were unfolded with buffer containing 6.0 M guanidinium hydrochloride (GdnHCl) for 4h at 4 °C. The unfolded LCs were then refolded in a buffer containing reduced and oxidized glutathione to assist in disulfide bond formation. The refolded proteins were subjected to anion exchange and size-exclusion chromatography steps for final purification. The level of protein purity was checked on 12% sodium dodecyl sulfate-polyacrylamide gel electrophoresis (SDS-PAGE) gels. The final protein concentration was measured using molecular weight and extinction coefficient of individual proteins. The purified proteins were stored at −20 °C for further use. Size exclusion coupled multi-angle light scattering (SEC-MALS) confirmed the dimeric assembly of purified proteins (**Figure S20**) which were used for SAXS and HDX-MS experiments mentioned below.

### Small-angle X-ray scattering (SAXS)

For SAXS analysis, H3 was diluted to 3.4 mg/mL, H7 was diluted to 3.4 mg/mL, H18 was diluted to 2.8 mg/mL, AL55 was diluted to 2.6 mg/mL, M7 was diluted to 3.6 mg/mL, in 20 mM TrisHCl, 150 mM NaCl, pH 8. H3, H7 and M7 batch data were collected at the P12 BioSAXS beamline of the EMBL Hamburg Synchrotron (52), while AL55 batch data and H18 and M10 SEC data were collected at the BM29 BioSAXS beamline of the ESRF, Grenoble (53). For SEC-SAXS, H18 and M10 were injected into a superdex 200 increase 10/300 GL column previously equilibrated in 20 mM TrisHCl, 150 mM NaCl, pH 8, at a concentration of 2.8 mg/mL and 6.7 mg/mL, respectively, see also **Figure S21** in the Supporting Information. SAXS data were processed using programs PRIMUS and GNOM within the ATSAS package (54). Data are deposited in the SASBDB (55) and available with accession codes SASDVL4, SASDVM4, SASDVN4, SASDVK4, SASDVP4, and SASDVQ4 (cf. **Dataset S1**).

### Molecular dynamics simulations

The available crystallographic structures of H3, H7 and M7 (PDB: 5mtl, 5muh and 5mvg, respectively (12)) were used as starting conformations, using Modeller to add missing residues (56). H18 and AL55 were modelled by homology modelling using SwissModel (57), while M10 was modelled using AF2 (58). Simulations were performed using GROMACS 2019 (59) and the PLUMED2 software (60–62), using AMBER-DES force field and TIP4P-D water (63,64). During in-vacuum minimization RMSD-restraints were imposed to enhance the symmetry between the two constant and the two variable domains. The systems were solvated in a periodic dodecahedron box, initially 1.2 nm larger than the protein in each direction, neutralized with Na and Cl ions to reach a salt concentration of 10 mM, then minimized and equilibrated at the temperature of 310 K and pressure of 1 atm using the Berendsen thermostat and barostat. Two independent 900 ns long plain MD simulations were run to generate reliable and independent starting conformations for the Metadynamics Metainference simulations (34,35,65). 30 conformations were extracted from each simulation and duplicated by inverting the two chains, to obtain 60 starting conformations symmetrically distributed with respect to chain inversion.

Metadynamics Metainference production simulations were run in duplicate using 60 replicas, each replica evolved for ∼1 μs (cf. **Table 2**). Simulations were performed in the NPT ensemble maintaining the temperature at 310 K with the Bussi thermostat (66) and the pressure of 1 atm with the Parrinello-Rahman barostat (67); the electrostatic was treated by using the particle mesh Ewald scheme with a short-range cut-off of 0.9 nm and van der Waals interaction cut-off was set to 0.9 nm. To reduce the computational cost, the hydrogen mass repartitioning scheme was used (68): the mass of heavy atoms was repartitioned into the bonded hydrogen atoms using the heavyh flag in the *pdb2gmx* tool, the LINCS algorithm was used to constraint all bonds, allowing to use a time step of 5 fs. In these simulations Parallel Bias Metadynamics (69) was used to enhance the sampling, combined with well-tempered metadynamics and the multiple-walker scheme, where Gaussians with an initial height of 1.0 kJ/mol were deposited every 0.5 ps, using a bias factor of 10. Five CVs were biased, including combinations of phi/psi dihedral angles of the linker regions (i.e. residues connecting variable and constant domains) in the two chains, combinations of chi dihedral angles of the linker regions in the two chains, combination of inter-domain contacts between the variable and the constant domains. The width of the Gaussians was 0.07, 0.12 and 120 for the combination of phi/psi, of chi dihedral angles and combination of contacts, respectively. Metainference was used to include SAXS restraints, using the hySAXS hybrid approach described in (36,37,70). A set of 13 representative SAXS intensities at different scattering angles, ranging between 0.015 AJJ and 0.25 AJJ and equally spaced, was used as restraints. These intensities were extracted from experimental data, after performing regularization with the Distance Distribution tool of Primus, based on Gnom (54). Metainference was applied every 5 steps, using a single Gaussian noise per data point and sampling a scaling factor between experimental and calculated SAXS intensities with a flat prior between 0.5 and 1.5. The aggregate sampling from the 60 replicas was reweighted using the final metadynamics bias to obtain a conformational ensemble where each conformation has an associated statistical weight (71). Convergence and error estimates were assessed by the inspection of the two replicated metadynamics metainference run. SAXS data were then recalculated using crysol (54). All relevant data are available on Zenodo (cf**. Dataset S2**).

### Hydrogen-deuterium mass exchange spectrometry (HDX-MS)

SYNAPT G2-HDMS system (Waters Corporation, USA) equipped with a LEAP robotic liquid handler was used to perform HDX-MS measurements in a fully automated mode as described previously (41,42,72,73). The data collection was carried out by a 20-fold dilution of H3, H7, AL55, and M10 proteins (100 µM) with the labeling buffer 1X phosphate buffer saline (PBS) prepared in D_2_O (pD 7.4) to trigger HDX for 0, 0.5, 1,10, 30, 120, and 240 min at 25 °C in technical triplicates. Each reaction was quenched by mixing the labeled protein with quench buffer (50 mM sodium phosphate, 250 mM TCEP, 3.0 M GdnHCl (pH 2)) in a 1:1 ratio at 0 °C. Online digestion was then performed using an immobilized pepsin digestion column (Waters Enzymate BEH Pepsin, 2.1 x 30 mm). The digested peptides were trapped using a C18 trapping column (Acquity BEH VanGuard 1.7 µm, 2.1 x 5.0 mm) and separated by a linear acetonitrile gradient of 5 to 40%. Protein Lynx Global Server (PLGS), and DynamX (Waters Corporation, USA) were used to identify the individual peptides, and subsequently, data processing using parameters: maximum peptide length of 25, the minimum intensity of 1000; minimum ion per amino acid of 0.1; maximum MH+ error of 5 ppm and a file threshold of three. A reference molecule [(Glu1)-fibrinopeptide B human (CAS No 103213-49-6, Merck, USA)] was used to lock mass with an expected molecular weight of 785.8426 Da. The obtained peptide coverage of H3, H7, AL55, and M10 were 98.6%, 92.5%, 98.6%, and 99.1%, respectively, with a redundancy > 4.0. Backbone amide groups exhibited a relative deuterium uptake (with respect to the zero exchange time data) of 0 to 30% within 4 h of exchange time. The relative deuterium uptake data at different HDX times were then used to generate heat maps for each amino acid. The obtained data was mapped on individual protein structures in a gradient of blue-white-red showing 0 to 30% of uptake, respectively (74). Red denotes the dynamic peptides while peptides colors in blue are rigid. The common peptide analysis between different model LCs was not performed in this analysis as they all generate different peptides both due to sequence heterogeneity in LCs especially in the VL domain and non-specific cleavage of pepsin enzyme used to generate peptides after deuterium exchanged for MS analysis. Therefore, all interpretation was done on individual relative exchange data.

## Acknowledgments

We acknowledge Martin A. Schroer and Dmitri Svergun at EMBL Hamburg, and Sonia Longhi at AFMB Marseille, for discussion and support on the SAXS data acquisition and analysis. We acknowledge PRACE for awarding us access to Piz Daint at CSCS, Switzerland. The authors acknowledge CINECA for an award under the ISCRA initiative for the availability of high-performance computing resources and support. This work was supported by the Italian Ministry of Research PRIN 2020 (20207XLJB2), by CARIPLO/TELETHON Foundations (GJC23044) and by Italian Ministry of Health (Ricerca Finalizzata, grant #GR-2018-12368387). SP acknowledges Fondazione Veronesi for a postdoctoral fellowship. STDH is supported by Academia Sinica intramural fund, an Academia Sinica Career Development Award from Academia Sinica (AS-CDA-109-L08), and the National Science and Technology Council (NSTC), Taiwan (NSTC113-2123-M-001-010-, 110-2113-M-001-050-MY3 and 110-2311-B-001-013-MY3). We also acknowledge Dr. Min-Feng Karen Hsu, and Mr. Yong-Sheng Wang for the protein quality controls, and Dr. Shu-Yu Lin, and Mr. Ming-Jie Tsai of the Academia Sinica Common Mass Spectrometry Facilities (AS-CFII-111-209), funded by the Academia Sinica Core Facility and Innovative Instrument Project, for supporting the HDX-MS experiments.

