## Supporting Information for "A conformational fingerprint for amyloidogenic light chains"

### Supporting Information for
A conformational fingerprint for amyloidogenic light chains.

Cristina Paissoni[1,a], Sarita Puri[1,a], Luca Broggini[2], Manoj K. Sriramoju[3], Martina Maritan[1], Rosaria Russo[1], Valentina Speranzini[1], Federico Ballabio[1], Mario Nuvolone[4], Giampaolo Merlini[4], Giovanni Palladini[4], Shang-Te Danny Hsu[3,5], Stefano Ricagno[1,2*], Carlo Camilloni[1*]

[1]Department of Bioscience, University of Milan, Milan, Italy.

[2]Institute of Molecular and Translational Cardiology, IRCCS, Policlinico San Donato, Milan, Italy.

[3]Institute of Biological Chemistry, Academia Sinica, Taipei, Taiwan.

[4]Department of Molecular Medicine, University of Pavia, Pavia, Italy; Amyloidosis Research and Treatment Center, Fondazione IRCCS Policlinico San Matteo, Pavia, Italy.

[5]Institute of Biomedical Sciences, National Taiwan University, Taipei, Taiwan; International Institute for Sustainability with Knotted Chiral Meta Matter (SKCM2), Hiroshima University, Higashi-Hiroshima, Japan.

[a]CP and SP contributed equally to this work and share the first authorship.

*Stefano Ricagno and Carlo Camilloni

**This PDF file includes:**

Figures S1 to S21
Tables S1 to S2
Legends for Movies S1 to S4
Legends for Datasets S1 to S2

**Other supporting materials for this manuscript include the following:**

Movies S1 to S4
Datasets S1 to S2

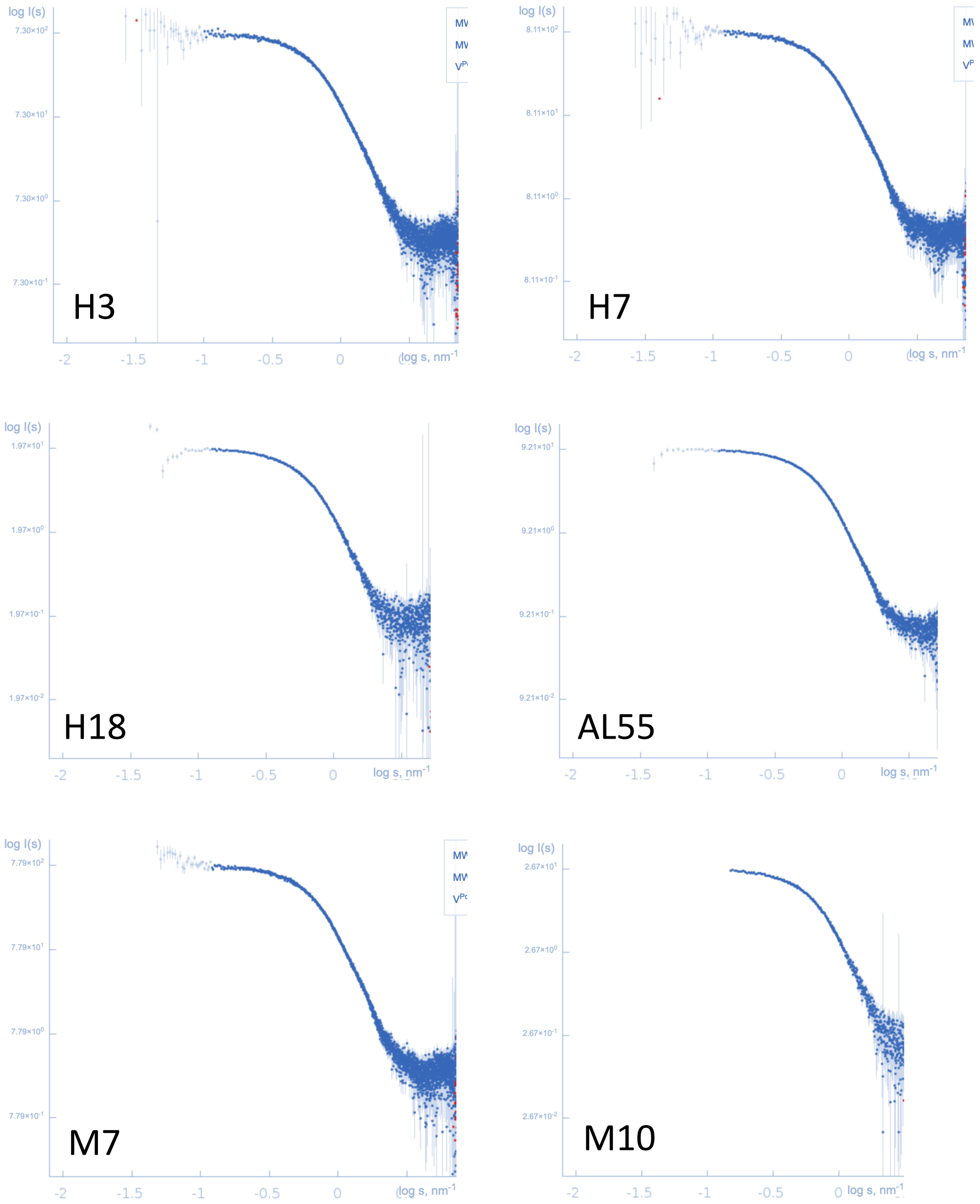


**Figure S1**: Log-Log plots representing the SAXS experimental curves for the six proteins.

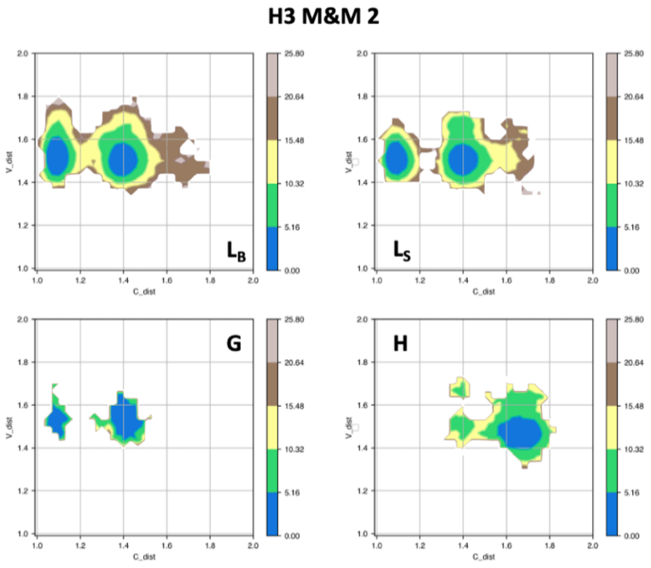


**Figure S2**: Free energy surfaces for the four substates identified in Figure 3 in the case of the second H3 metainference simulation. The x-axis shows the distance between the centers of mass of the constant domains, while the y-axis shows the distance between the centers of mass of the variable domains. The free energy is shown with color and isolines every $2k_BT$ corresponding to 5.16 kJ/mol.

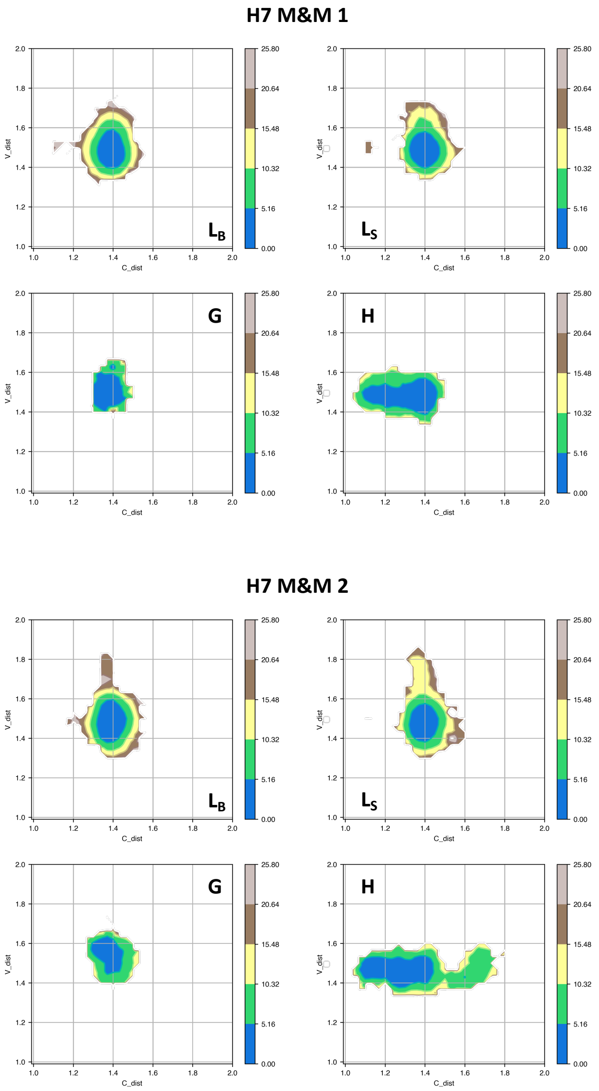


**Figure S3**: Free energy surfaces for the four substates identified in Figure 3 in the case of the first (top) and second (bottom) H7 metainference simulation. The x-axis shows the distance between the centers of mass of the constant domains, while the y-axis shows the distance between the centers of mass of the variable domains. The free energy is shown with color and isolines every $2k_BT$ corresponding to 5.16 kJ/mol.

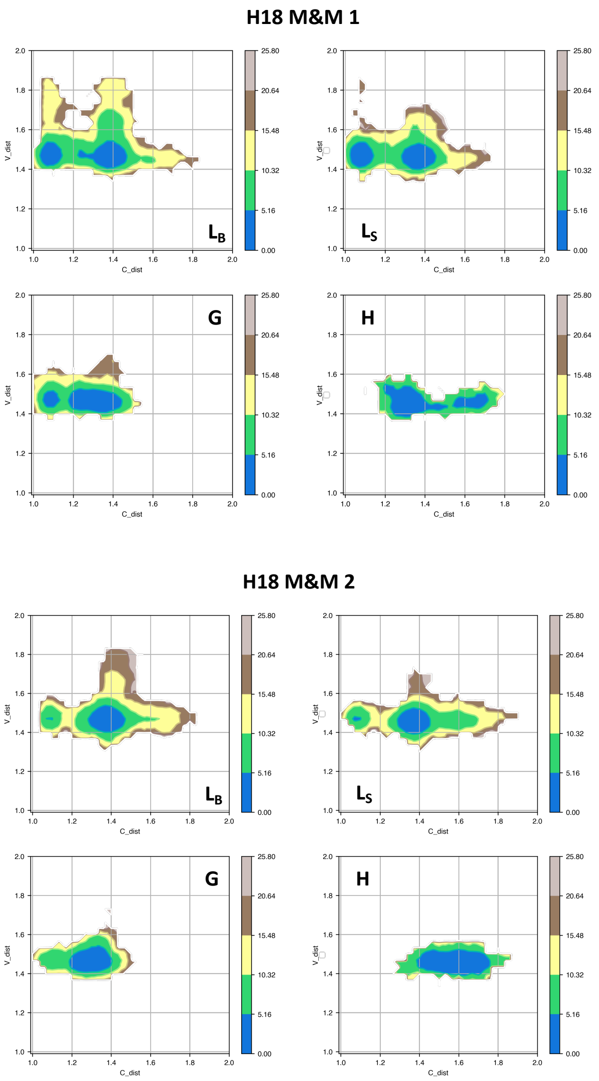


**Figure S4**: Free energy surfaces for the four substates identified in Figure 3 in the case of the first (top) and second (bottom) H18 metainference simulation. The x-axis shows the distance between the centers of mass of the constant domains, while the y-axis shows the distance between the centers of mass of the variable domains. The free energy is shown with color and isolines every $2k_BT$ corresponding to 5.16 kJ/mol.

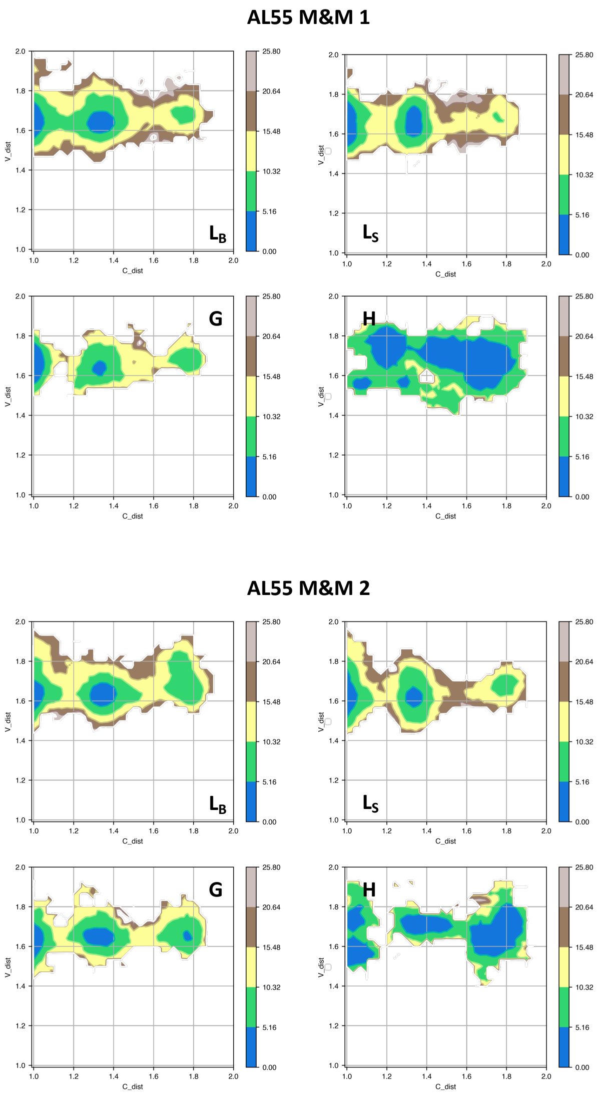


**Figure S5:** Free energy surfaces for the four substates identified in Figure 3 in the case of the first (top) and second (bottom) AL55 metainference simulation. The x-axis shows the distance between the centers of mass of the constant domains, while the y-axis shows the distance between the centers of mass of the variable domains. The free energy is shown with color and isolines every $2k_BT$ corresponding to 5.16 kJ/mol.

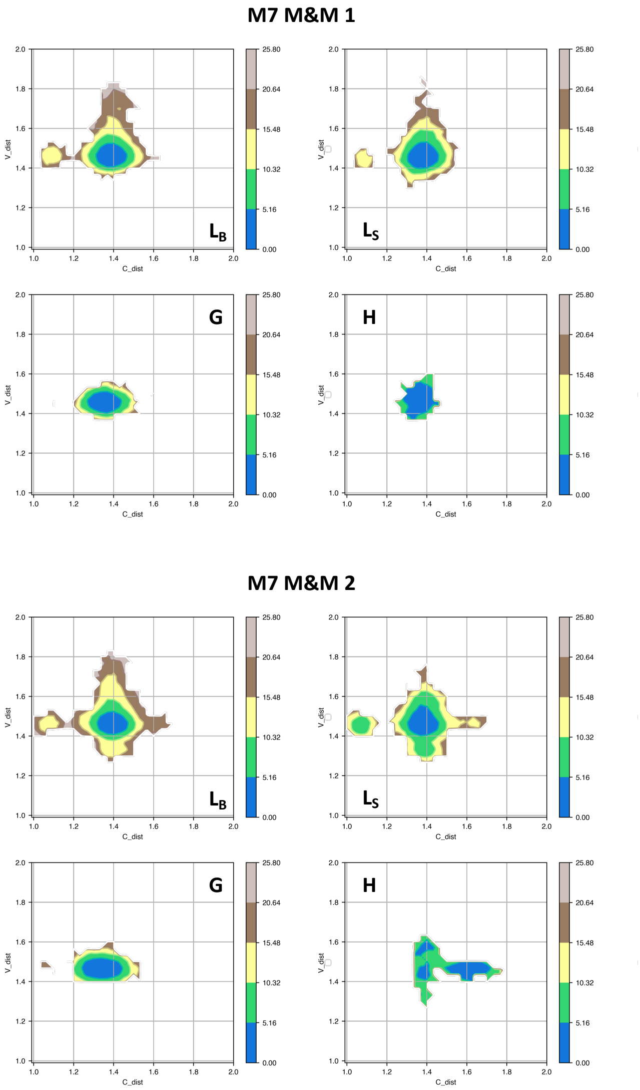


**Figure S6:** Free energy surfaces for the four substates identified in Figure 3 in the case of the first (top) and second (bottom) M7 metainference simulation. The x-axis shows the distance between the centers of mass of the constant domains, while the y-axis shows the distance between the centers of mass of the variable domains. The free energy is shown with color and isolines every $2k_BT$ corresponding to 5.16 kJ/mol.

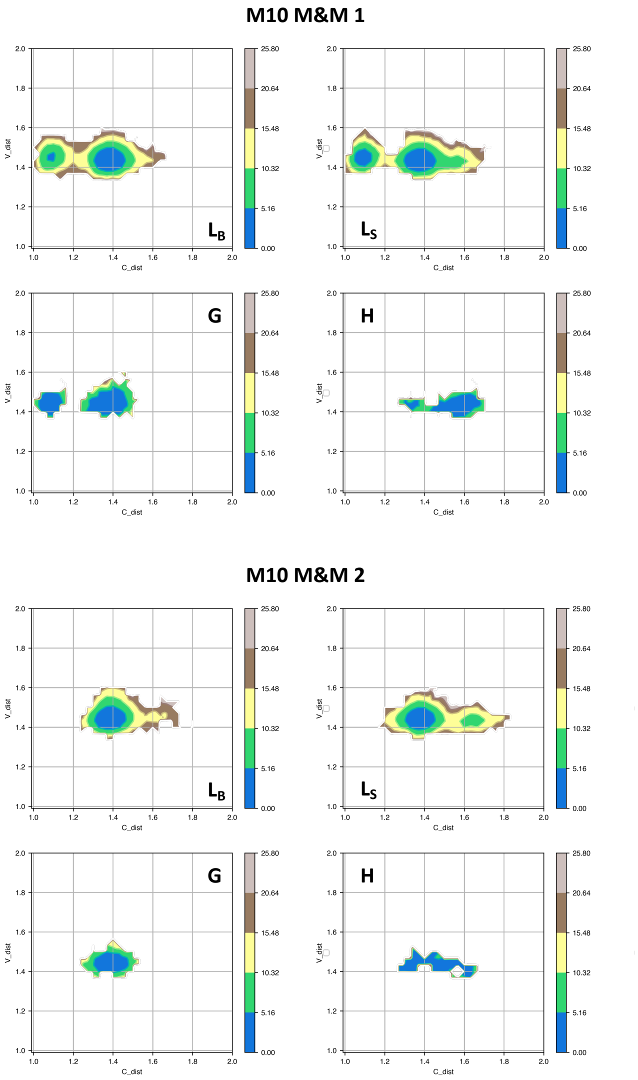


**Figure S7:** Free energy surfaces for the four substates identified in Figure 3 in the case of the first (top) and second (bottom) M10 metainference simulation. The x-axis shows the distance between the centers of mass of the constant domains, while the y-axis shows the distance between the centers of mass of the variable domains. The free energy is shown with color and isolines every $2k_BT$ corresponding to 5.16 kJ/mol.

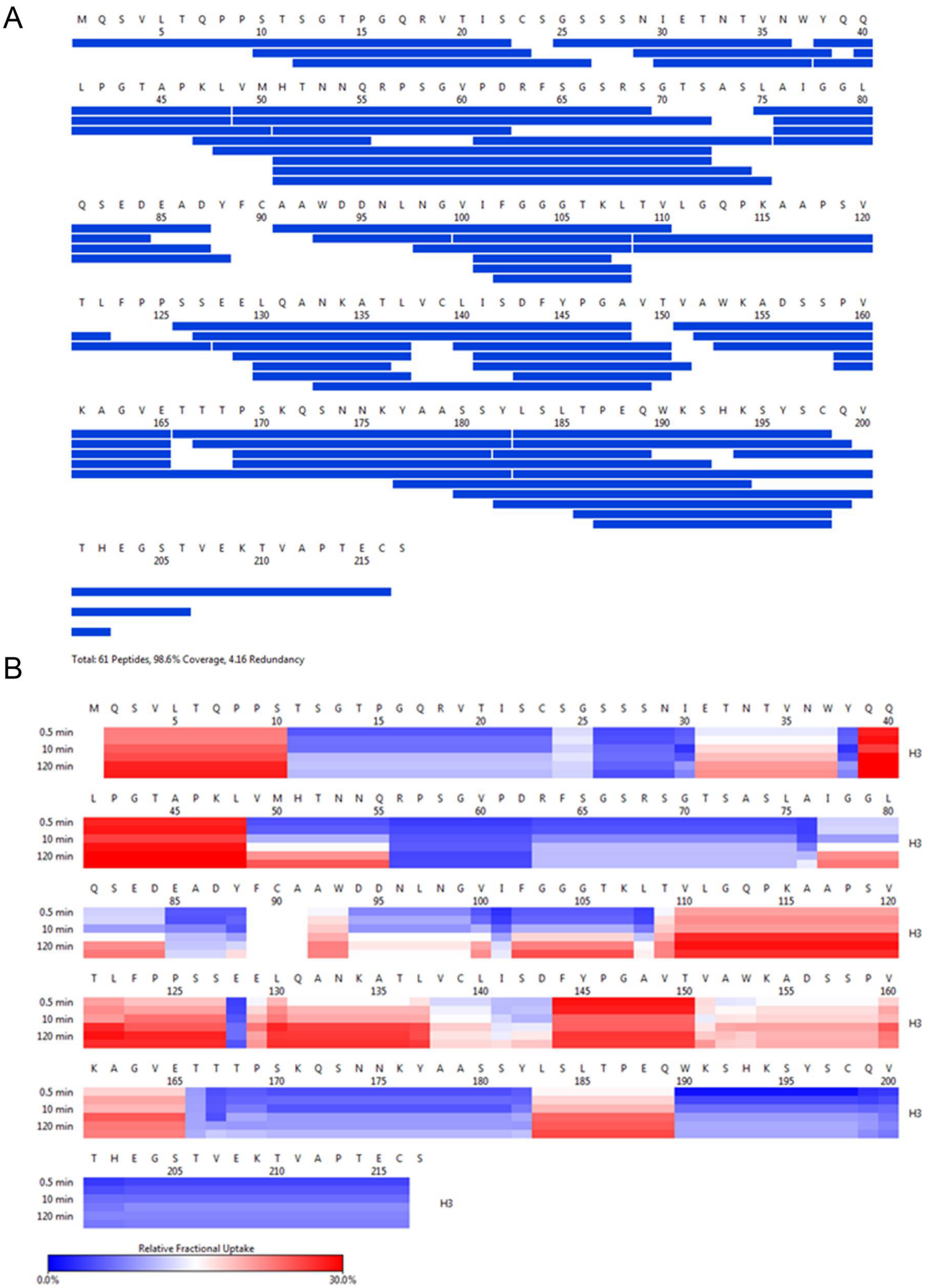


**Figure S8**: (A) Peptide coverage map of protein H3. The total number of peptides is 61 with a coverage of 98.6% and a redundancy of 4.16. (B) As shown on the left, heat map as a function of HDX-time at different time points. The relative deuterium uptake is color-coded from blue-to-white-to-red for 0 to 30% as indicated by the scale bar below.

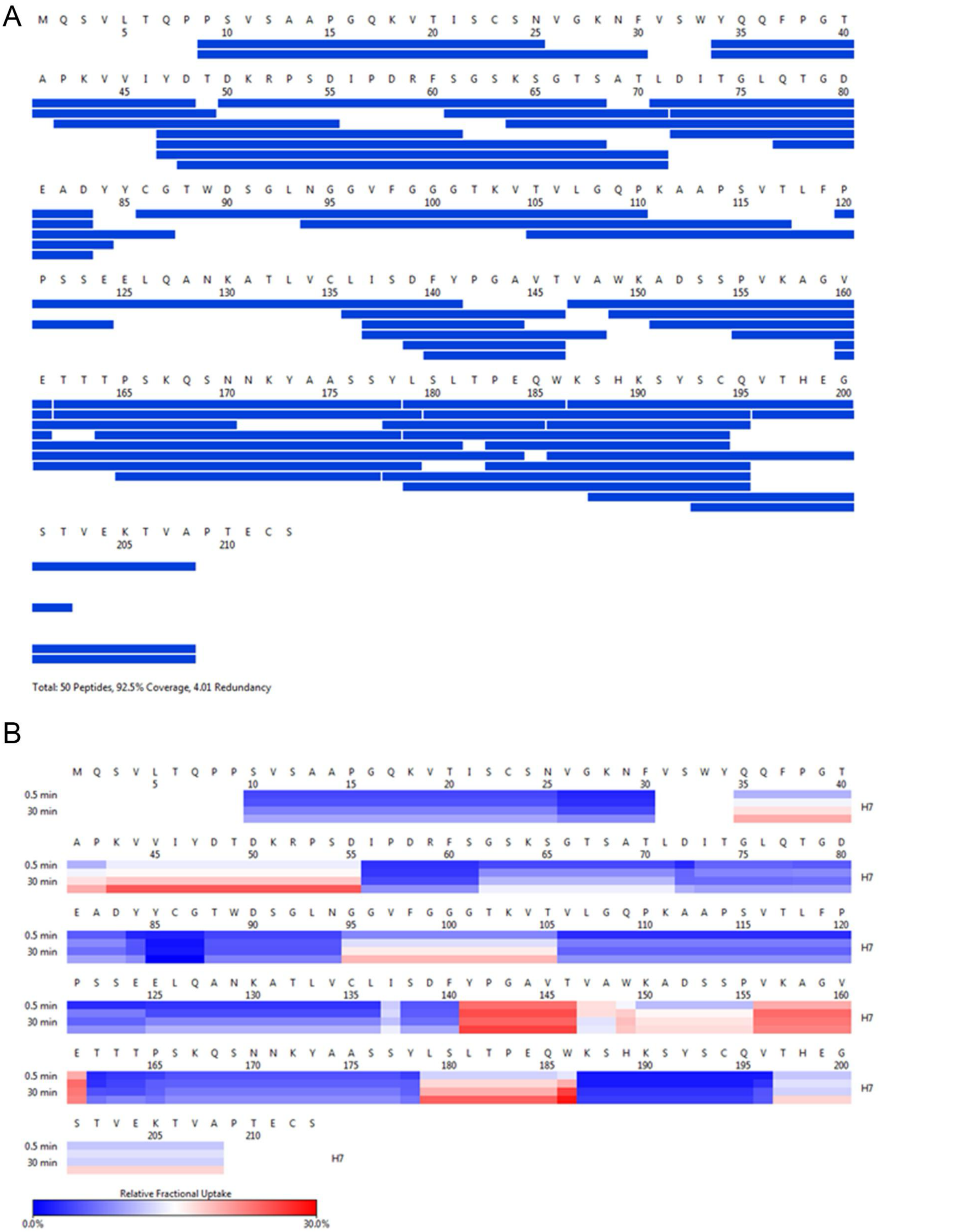


**Figure S9**: (A) Peptide coverage map of protein H7. The total number of peptides is 50 with a coverage of 92.5% and a redundancy of 4.01. (B) Heat map as a function of HDX-time at different time points as shown on the left. The relative deuterium uptake is color-coded from blue-to-white-to-red for 0 to 30% as shown by the scale bar below.

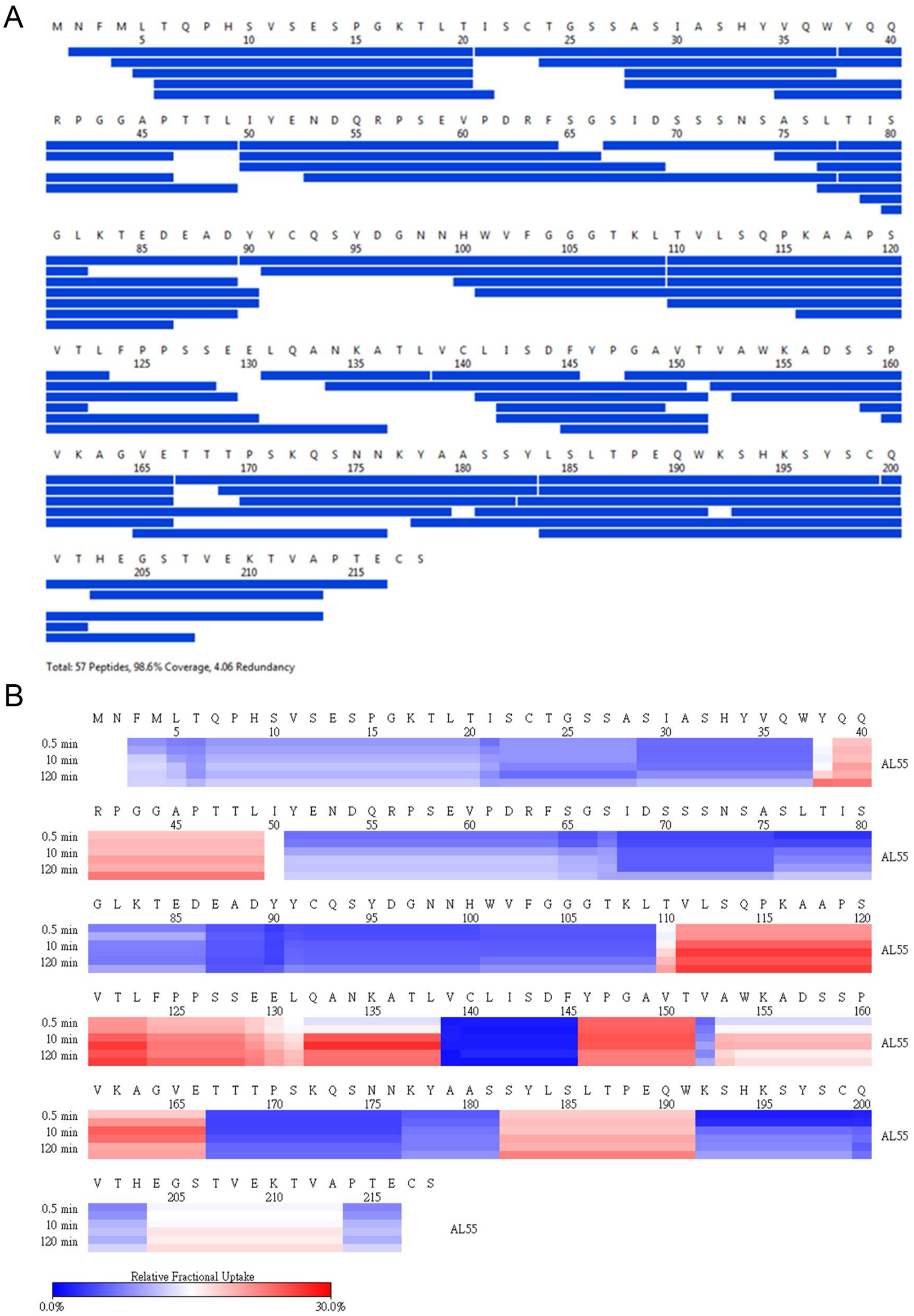


**Figure S10**: (A) Peptide coverage map of protein AL55. The total number of peptides is 57 with a coverage of 98.6% and a redundancy of 4.06. (B) Heat map as a function of HDX-time at different time points as shown on the left. The relative deuterium uptake is color-coded from blue-to-white-to-red for 0 to 30% as shown by the scale bar below.

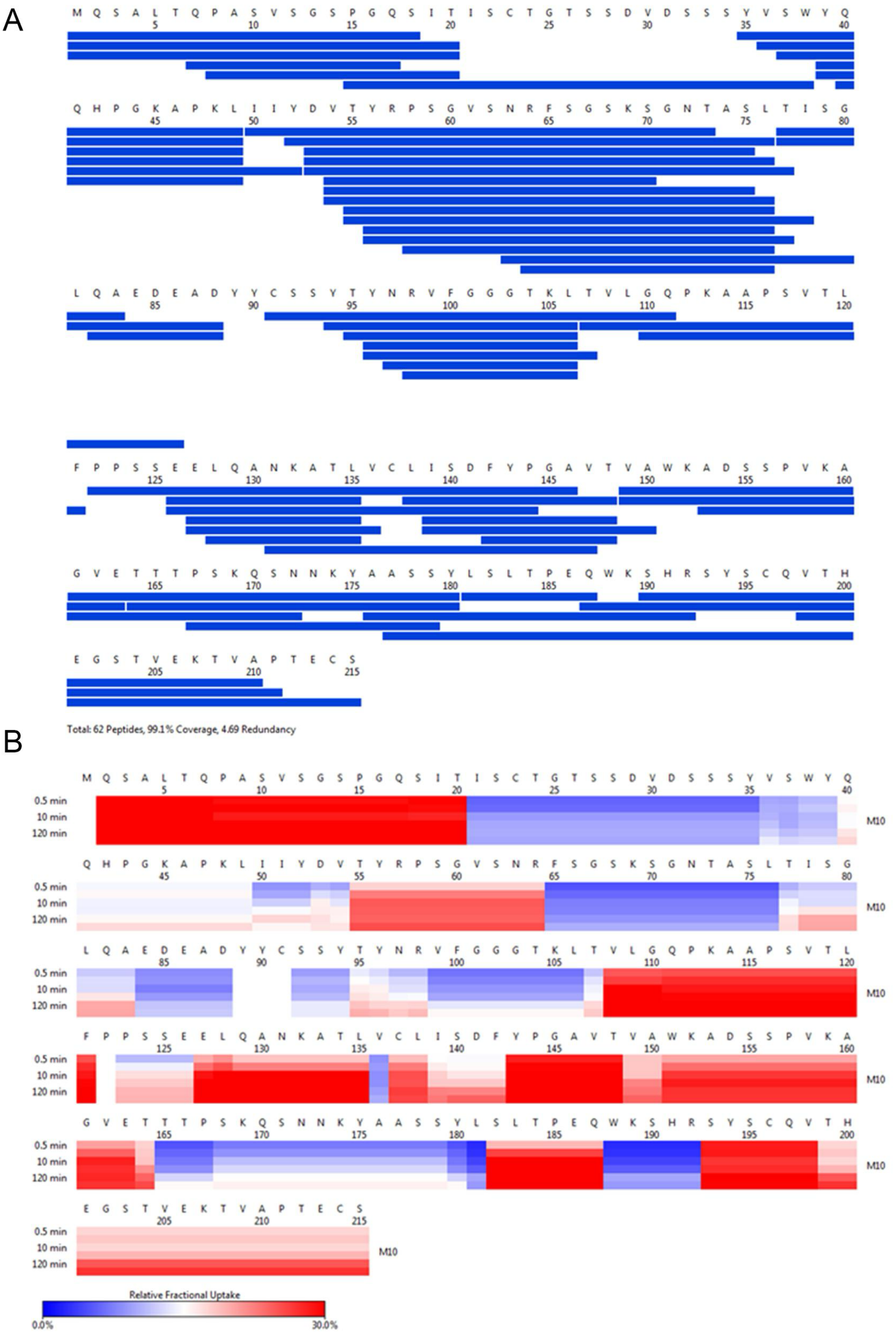


**Figure S11**: (A) Peptide coverage map of protein M10. The total number of peptides is 62 with a coverage of 99.1% and a redundancy of 4.69. (B) As shown on the left, heat map as a function of HDX-time at different time points. The relative deuterium uptake is color-coded from blue-to-white-to-red for 0 to 30% as indicated by the scale bar below.

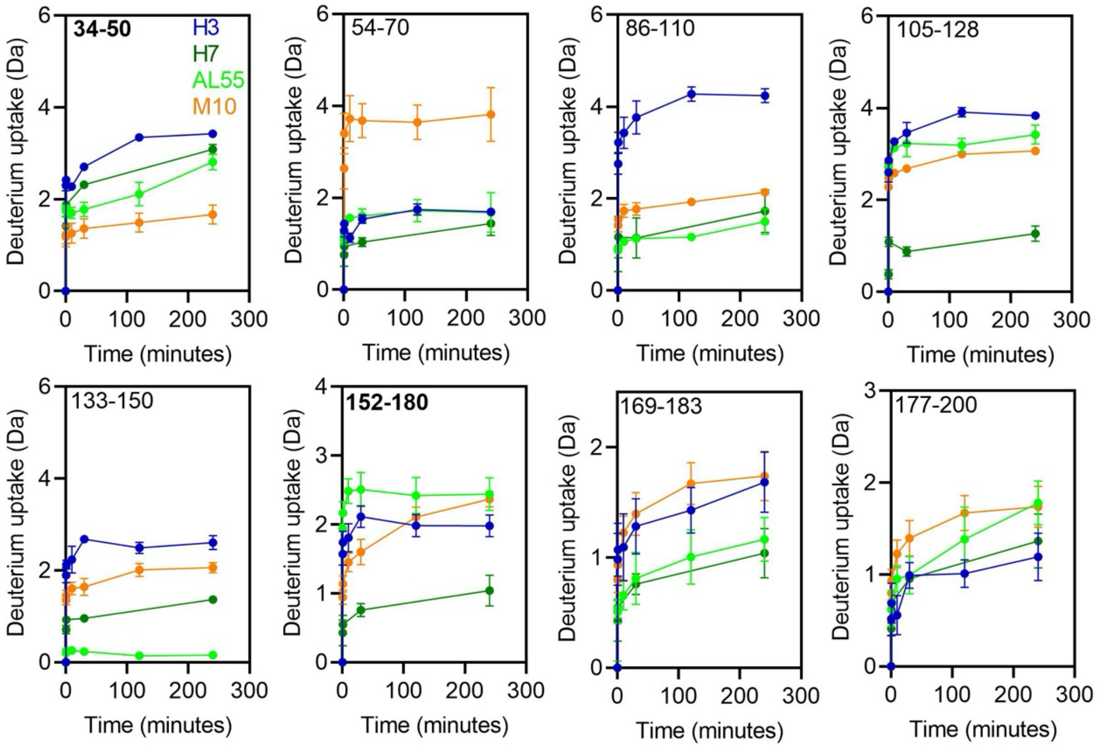


**Figure S12**: HDX kinetics of deuterium uptake at each time points from 0 to 240 min for selected peptides. The color corresponding to each protein H3 (blue), H7 (green), AL55 (light green), and M10 (orange) are shown in first panel. The residue numbers corresponding to the individual peptides are shown on the upper left corner of the individual panels. Panel for peptides containing residues 34-50 and 152-180 highlighted in bold are of our interest for this study.

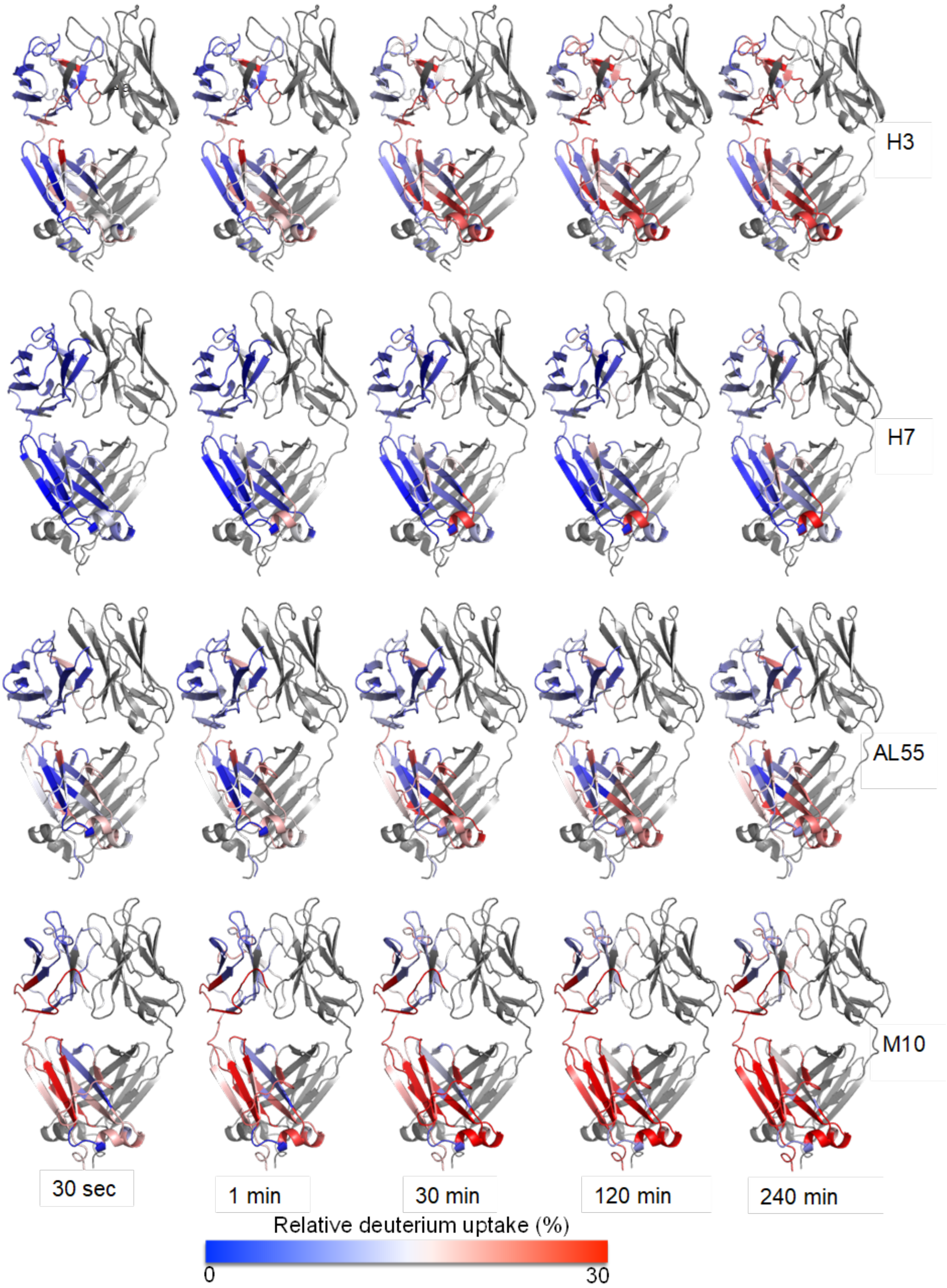


**Figure S13**: **Structural mapping of relative deuterium uptake on chain A of each dimeric LC**: Relative deuterium uptake of H3, H7, AL55, and M10 at different time points of 0.5 to 240 min on a scale of 0 to 30% uptake. The color gradient is from blue-white-red. Blue means no exchange and red means an exchange of 30%. The protein name is written on right hand side of each panel. Chain B is colored in gray.

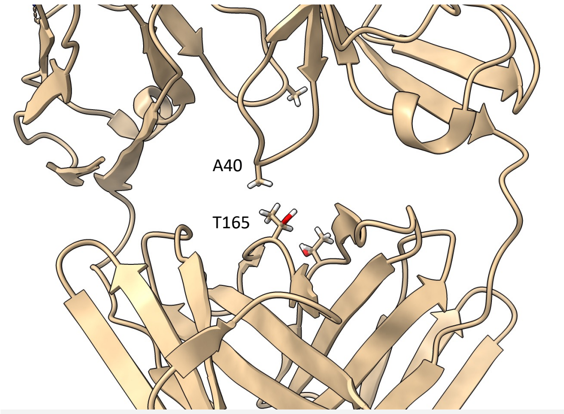


```
M7        SSELTQDPAVSVALGQTVKITCQGDSLRMYYASWYQQKPAQAPVLVIYAEKNRPSGIPDR
H18       SSQLTQDPAVSVALGQIVTITCQGDSLRTYYASWYQQKPGQAPVLVIYNQDHRPSGIPDR
germline: SSELTQDPAVSVALGQTVRITCQGDSLRSYYASWYQQKPGQAPVLVIYGKNNRPSGIPDR
          **:************* * ********* **********.******** :.:********
                            20                  40                  60

M7        FSASSSGSTASLTITGAQAEDEADYYCNSRDNSGDHLVFGGGTKLTVLGQPKAAPSVTLF
H18       FSGSSSGNTASLTIAGAQANDEADYYCNSRDSSGNLVLFGGGTKLTVLGQPKAAPSVTLF
germline: FSGSSSGNTASLTITGAQAEDEADYYCNSRDSSGNLVLFGGGTKLTVLGQPKAAPSVTLF
          **.****.******:****:***********.**: ::**********************
                            80                 100                 120

M7        PPSSEELQANKATLVCLISDFYPGAVTVAWKADSSPVKAGVETTTPSKQSNNKYAASSYL
H18       PPSSEELQANKATLVCLISDFYPGAVTVAWKADSSPVKAGVETTTPSKQSNNKYAASSYL
germline: PPSSEELQANKATLVCLISDFYPGAVTVAWKADSSPVKAGVETTTPSKQSNNKYAASSYL
          ************************************************************
                           140                 160                180

M7        SLTPEQWKSHRSYSCQVTHEGSTVEKTVAPTECS
H18       SLTPEQWKSHRSYSCQVTHEGSTVEKTVAPTECS
germline: SLTPEQWKSHRSYSCQVTHEGSTVEKTVAPTECS
          **********************************
                           200
```

**Figure S13**. Zoom in on the crystal structure of M7 to show the hydrophobic contact between A40 in the VL and T165 in the CL. Below is reported the multiple sequence alignment for M7, H18 and their germline sequence (*IGLV3-19*01* for the VL and *IGLC2*02* for the CL).

```
H3                 1 QSVLTQPPSTSGTPGQRVTISCSGSSSNIETNTVNWYQQLPGTAPKLVMH    50
                     |||||||||.|||||||||||||||||||.:||||||||||||||||:::
IGLV1-44           1 QSVLTQPPSASGTPGQRVTISCSGSSSNIGSNTVNWYQQLPGTAPKLLIY    50

H3                51 TNNQRPSGVPDRFSGSRSGTSASLAIGGLQSEDEADYFCAAWDDNLNGVI   100
                     :|||||||||||||||:|||||||||.||||||||||:||||||:|||||
IGLV1-44          51 SNNQRPSGVPDRFSGSKSGTSASLAISGLQSEDEADYYCAAWDDSLNGVI   100

H3               101 FGGGTKLTVLGQPKAAPSVTLFPPSSEELQANKATLVCLISDFYPGAVTV   150
                     ||||||||||||||||||||||||||||||||||||||||||||||||||
IGLV1-44/C3*03   101 FGGGTKLTVLGQPKAAPSVTLFPPSSEELQANKATLVCLISDFYPGAVTV   150

H3               151 AWKADSSPVKAGVETTTPSKQSNNKYAASSYLSLTPEQWKSHKSYSCQVT   200
                     ||||||||||||||||||||||||||||||||||||||||||||||||||
IGLC3*03         151 AWKADSSPVKAGVETTTPSKQSNNKYAASSYLSLTPEQWKSHKSYSCQVT   200

H3               201 HEGSTVEKTVAPTECS    216
                     ||||||||||||||||
IGLC3*03         201 HEGSTVEKTVAPTECS    216
```

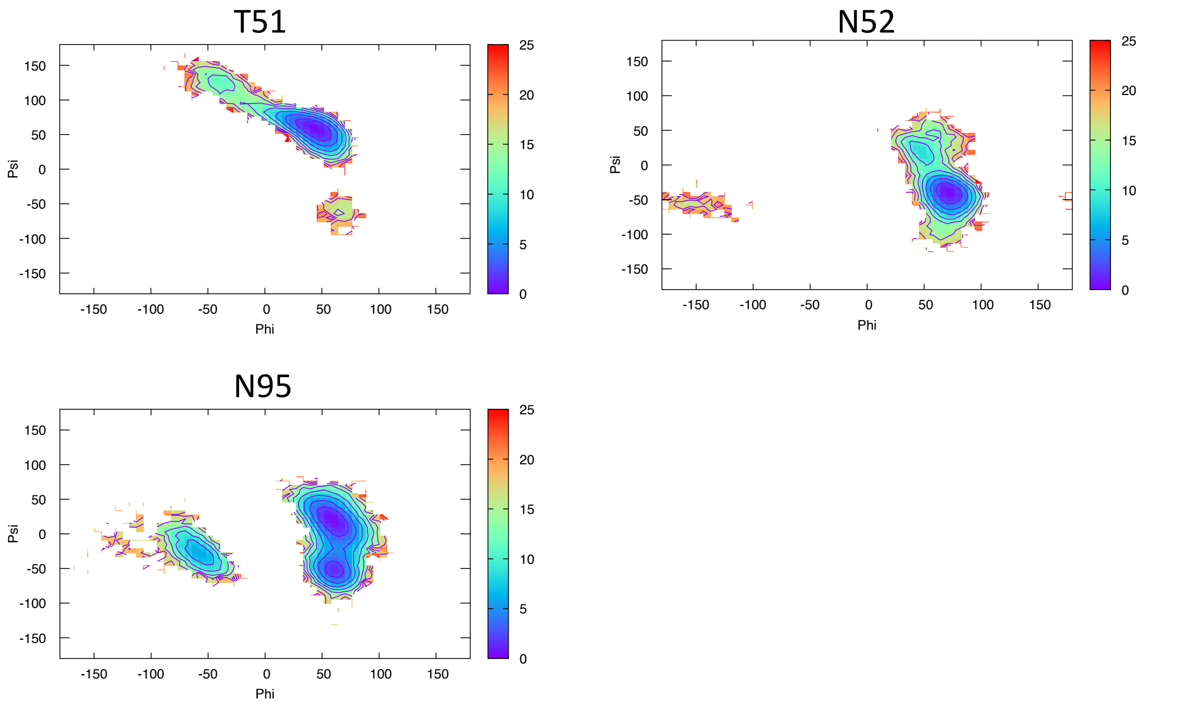


**Figure S14:** Pairwise sequence alignment between H3 and its corresponding germline as identified by igBLAST using the IGMT databases. The three CDRs and the linker region are highlighted in light blue and orange, respectively. The red circles indicate residues for which the left alpha is the most populated region in the Ramachandran plot. FES (in kJ/mol) representing the Ramachandran plot for the indicated residues are reported in the bottom panels.

```
H7               1  QSVLTQPPSVSAAPGQKVTISC----SNVGKNFVSWYQQFPGTAPKVVIY   46
                    ||||||||||||||||||||||    ||:|.|:||||||.||||||::||
IGLV1-51*01      1  QSVLTQPPSVSAAPGQKVTISCSGSSSNIGNNYVSWYQQLPGTAPKLLIY   50

H7              47  DTDKRPSDIPDRFSGSKSGTSATLDITGLQTGDEADYYCGTWDSGLNGGV   96
                    |.:||||.||||||||||||||||.|||||||||||||||||||.|:.||
IGLV1-51*01     51  DNNKRPSGIPDRFSGSKSGTSATLGITGLQTGDEADYYCGTWDSSLSAGV  100

H7              97  FGGGTKVTVLGQPKAAPSVTLFPPSSEELQANKATLVCLISDFYPGAVTV  146
                    ||||||||||||||||||||||||||||||||||||||||||||||||||
IGLV1-51/C3*03 101  FGGGTKVTVLGQPKAAPSVTLFPPSSEELQANKATLVCLISDFYPGAVTV  150

H7             147  AWKADSSPVKAGVETTTPSKQSNNKYAASSYLSLTPEQWKSHKSYSCQVT  196
                    ||||||||||||||||||||||||||||||||||||||||||||||||||
IGLC3*03 .      151  AWKADSSPVKAGVETTTPSKQSNNKYAASSYLSLTPEQWKSHKSYSCQVT  200

H7             197  HEGSTVEKTVAPTECS   212
                    ||||||||||||||||
IGLC3*03       201  HEGSTVEKTVAPTECS   216
```

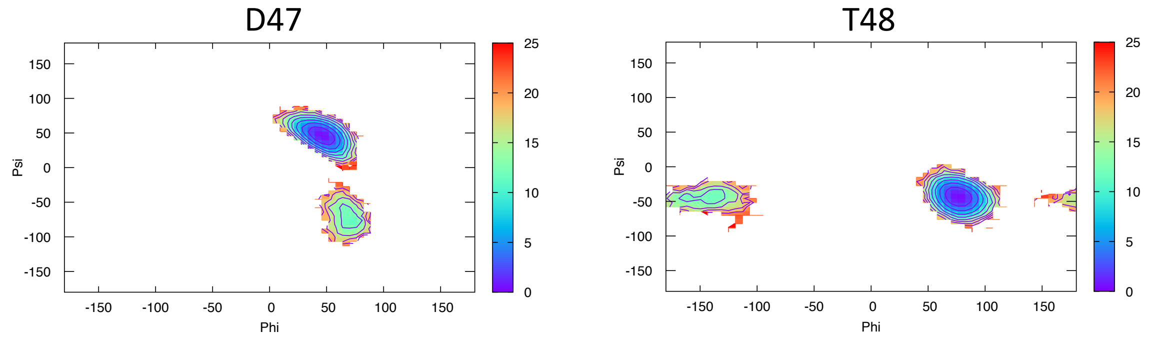


**Figure S15:** Pairwise sequence alignment between H7 and its corresponding germline as identified by igBLAST using the IGMT databases. The three CDRs and the linker region are highlighted in light blue and orange, respectively. The red circles indicate residues for which the left alpha is the most populated region in the Ramachandran plot. FES (in kJ/mol) representing the Ramachandran plot for the indicated residues are reported in the bottom panels.

```
H18              1 SSQLTQDPAVSVALGQIVTITCQGDSLRTYYASWYQQKPGQAPVLVIYNQ     50
                   ||:|||||||||||||.|.|||||||||:|||||||||||||||||||.:
IGLV3-19*01      1 SSELTQDPAVSVALGQTVRITCQGDSLRSYYASWYQQKPGQAPVLVIYGK     50

H18             51 DHRPSGIPDRFSGSSSGNTASLTIAGAQANDEADYYCNSRDSSGNLVLFG    100
                   ::||||||||||||||||||||||.||||.||||||||||||||||||||
IGLV3-19*01     51 NNRPSGIPDRFSGSSSGNTASLTITGAQAEDEADYYCNSRDSSGNLVLFG    100

H18            101 GGTKLTVLGQPKAAPSVTLFPPSSEELQANKATLVCLISDFYPGAVTVAW    150
                   ||||||||||||||||||||||||||||||||||||||||||||||||||
IGLV3-19/C2*02 101 GGTKLTVLGQPKAAPSVTLFPPSSEELQANKATLVCLISDFYPGAVTVAW    150

H18            151 KADSSPVKAGVETTTPSKQSNNKYAASSYLSLTPEQWKSHRSYSCQVTHE    200
                   ||||||||||||||||||||||||||||||||||||||||||||||||||
IGLC2*02       151 KADSSPVKAGVETTTPSKQSNNKYAASSYLSLTPEQWKSHRSYSCQVTHE    200

H18            201 GSTVEKTVAPTECS    214
                   ||||||||||||||
IGLC2*02       201 GSTVEKTVAPTECS    214
```

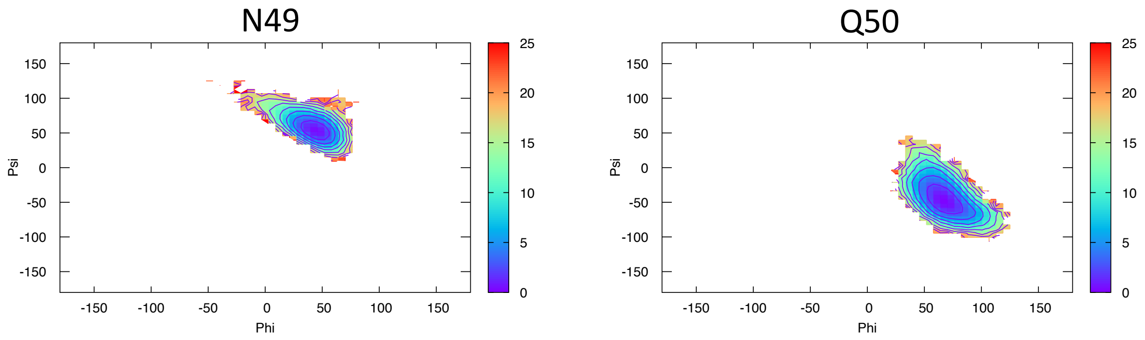


**Figure S16:** Pairwise sequence alignment between H18 and its corresponding germline as identified by igBLAST using the IGMT databases. The three CDRs and the linker region are highlighted in light blue and orange, respectively. The red circles indicate residues for which the left alpha is the most populated region in the Ramachandran plot. FES (in kJ/mol) representing the Ramachandran plot for the indicated residues are reported in the bottom panels.

```
AL55              1  NFMLTQPHSVSESPGKTLTISCTGSSASIASHYVQWYQQRPGGAPTTLIY   50
                     |||||||||||||||||:||||||||.||||:||||||||||.||||:||
IGLV6-57*02       1  NFMLTQPHSVSESPGKTVTISCTGSSGSIASNYVQWYQQRPGSAPTTVIY   50

AL55             51  ENDQRPSEVPDRFSGSIDSSSNSASLTISGLKTEDEADYYCQSYDGNNHW  100
                     |::||||.|||||||||||||||||||||||||||||||||||||.:|||
IGLV6-57*02      51  EDNQRPSGVPDRFSGSIDSSSNSASLTISGLKTEDEADYYCQSYDSSNHW  100

AL55            101  VFGGGTKLTVLSQPKAAPSVTLFPPSSEELQANKATLVCLISDFYPGAVT  150
                     ||||||||||||||||||||||||||||||||||||||||||||||||||
IGLV6-57/C3*03  101  VFGGGTKLTVLSQPKAAPSVTLFPPSSEELQANKATLVCLISDFYPGAVT  150

AL55            151  VAWKADSSPVKAGVETTTPSKQSNNKYAASSYLSLTPEQWKSHKSYSCQV  200
                     ||||||||||||||||||||||||||||||||||||||||||||||||||
IGLC3*03        151  VAWKADSSPVKAGVETTTPSKQSNNKYAASSYLSLTPEQWKSHKSYSCQV  200

AL55            201  THEGSTVEKTVAPTECS  217
                     |||||||||||||||||
IGLC3*03        201  THEGSTVEKTVAPTECS  217
```

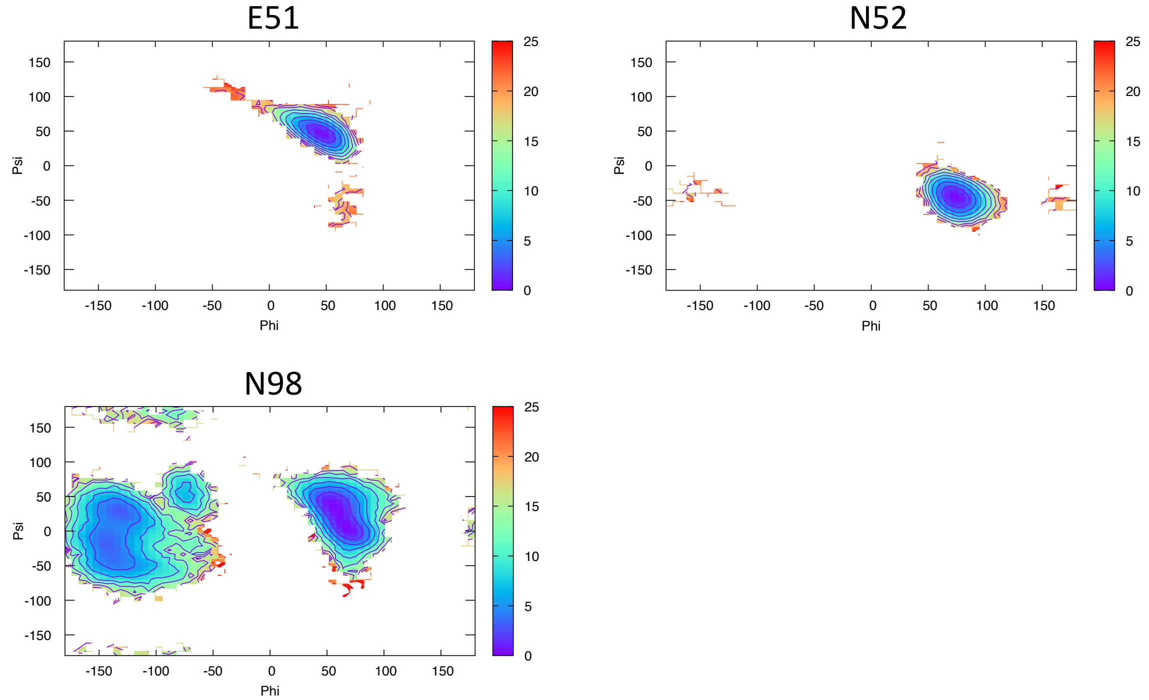


**Figure S17:** Pairwise sequence alignment between AL55 and its corresponding germline as identified by igBLAST using the IGMT databases. The three CDRs and the linker region are highlighted in light blue and orange, respectively. The red circles indicate residues for which the left alpha is the most populated region in the Ramachandran plot. FES (in kJ/mol) representing the Ramachandran plot for the indicated residues are reported in the bottom panels.

```
M7                1 SSELTQDPAVSVALGQTVKITCQGDSLRMYYASWYQQKPAQAPVLVIYAE    50
                    ||||||||||||||||||:|||||||||.||||||||||.||||||||.:
IGLV3-19*01       1 SSELTQDPAVSVALGQTVRITCQGDSLRSYYASWYQQKPGQAPVLVIYGK    50

M7               51 KNRPSGIPDRFSASSSGSTASLTITGAQAEDEADYYCNSRDNSGDHLVFG   100
                    .|||||||||||.||||:|||||||||||||||||||||||:||:|||||
IGLV3-19*01      51 NNRPSGIPDRFSGSSSGNTASLTITGAQAEDEADYYCNSRDSSGNHLVFG   100

M7              101 GGTKLTVLGQPKAAPSVTLFPPSSEELQANKATLVCLISDFYPGAVTVAW   150
                    ||||||||||||||||||||||||||||||||||||||||||||||||||
IGLV3-19/C2*02  101 GGTKLTVLGQPKAAPSVTLFPPSSEELQANKATLVCLISDFYPGAVTVAW   150

M7              151 KADSSPVKAGVETTTPSKQSNNKYAASSYLSLTPEQWKSHRSYSCQVTHE   200
                    ||||||||||||||||||||||||||||||||||||||||||||||||||
IGLC2*02        151 KADSSPVKAGVETTTPSKQSNNKYAASSYLSLTPEQWKSHRSYSCQVTHE   200

M7              201 GSTVEKTVAPTECS    214
                    ||||||||||||||
IGLC2*02        201 GSTVEKTVAPTECS    214
```

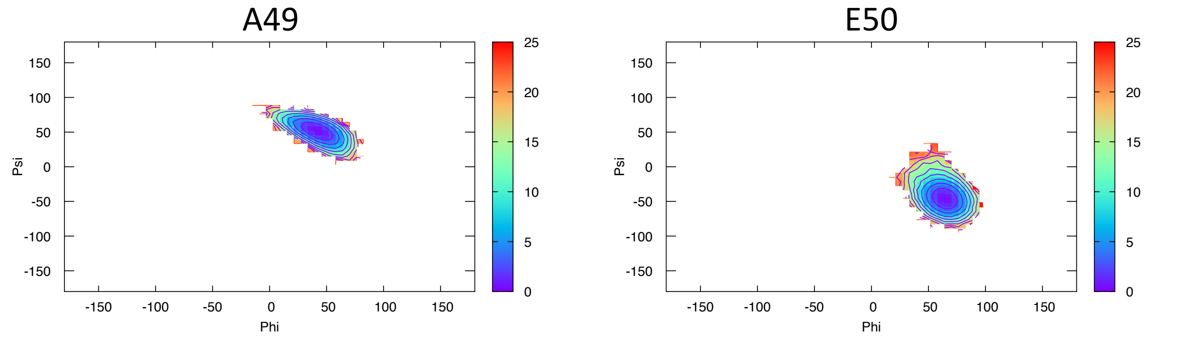


**Figure S18:** Pairwise sequence alignment between M7 and its corresponding germline as identified by igBLAST using the IGMT databases. The three CDRs and the linker region are highlighted in light blue and orange, respectively. The red circles indicate residues for which the left alpha is the most populated region in the Ramachandran plot. FES (in kJ/mol) representing the Ramachandran plot for the indicated residues are reported in the bottom panels.

```
M10                 1 QSALTQPASVSGSPGQSITISCTGTSSDVDSSSYVSWYQQHPGKAPKLII    50
                      |||||||||||||||||||||||||||||...:|||||||||||||||:|
IGLV2-14*01         1 QSALTQPASVSGSPGQSITISCTGTSSDVGGYNYVSWYQQHPGKAPKLMI    50

M10                51 YDVTYRPSGVSNRFSGSKSGNTASLTISGLQAEDEADYYCSSYTYNRVFG   100
                      |:|:.|||||||||||||||||||||||||||||||||||||||.:||||
IGLV2-14*01        51 YEVSNRPSGVSNRFSGSKSGNTASLTISGLQAEDEADYYCSSYTSSRVFG   100

M10               101 GGTKLTVLGQPKAAPSVTLFPPSSEELQANKATLVCLISDFYPGAVTVAW   150
                      ||||||||||||||||||||||||||||||||||||||||||||||||||
IGLV2-14/C3*04    101 GGTKLTVLGQPKAAPSVTLFPPSSEELQANKATLVCLISDFYPGAVTVAW   150

M10               151 KADSSPVKAGVETTTPSKQSNNKYAASSYLSLTPEQWKSHRSYSCQVTHE   200
                      ||||||||||||||||||||||||||||||||||||||||||||||||||
IGLC3*04          151 KADSSPVKAGVETTTPSKQSNNKYAASSYLSLTPEQWKSHRSYSCQVTHE   200

M10               201 GSTVEKTVAPTECSc    215
                      ||||||||||||||
IGLC3*04          201 GSTVEKTVAPTECS-    214
```

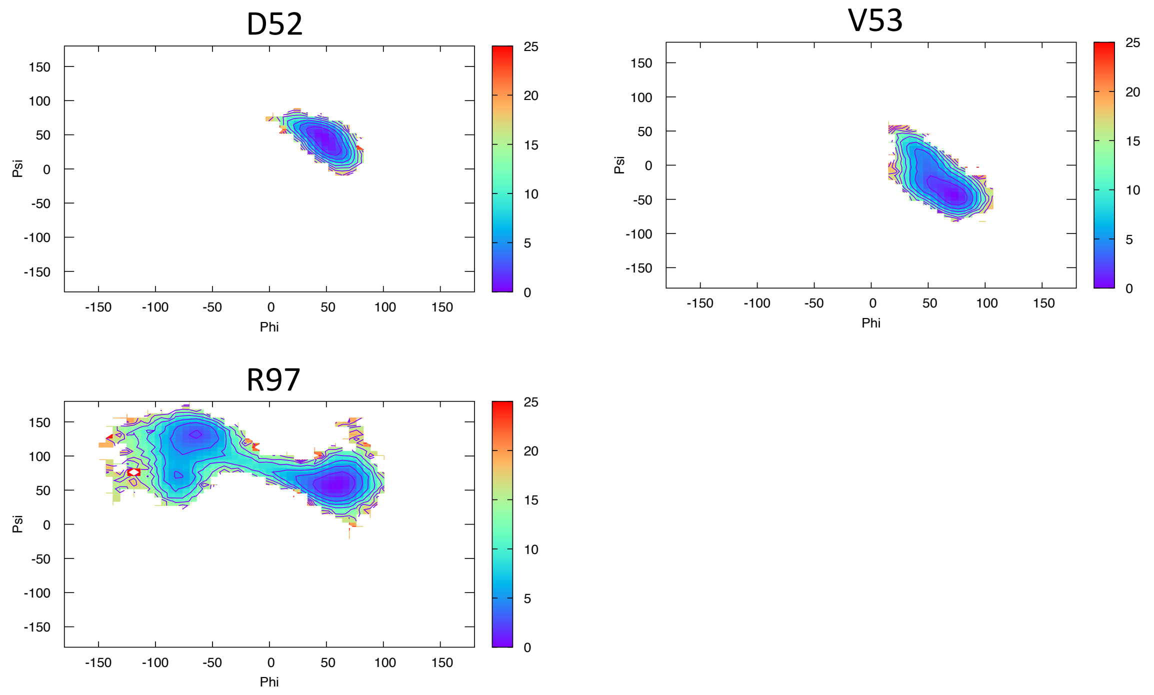


**Figure S19:** Pairwise sequence alignment between M10 and its corresponding germline as identified by igBLAST using the IGMT databases. The three CDRs and the linker region are highlighted in light blue and orange, respectively. The red circles indicate residues for which the left alpha is the most populated region in the Ramachandran plot. FES (in kJ/mol) representing the Ramachandran plot for the indicated residues are reported in the bottom panels.

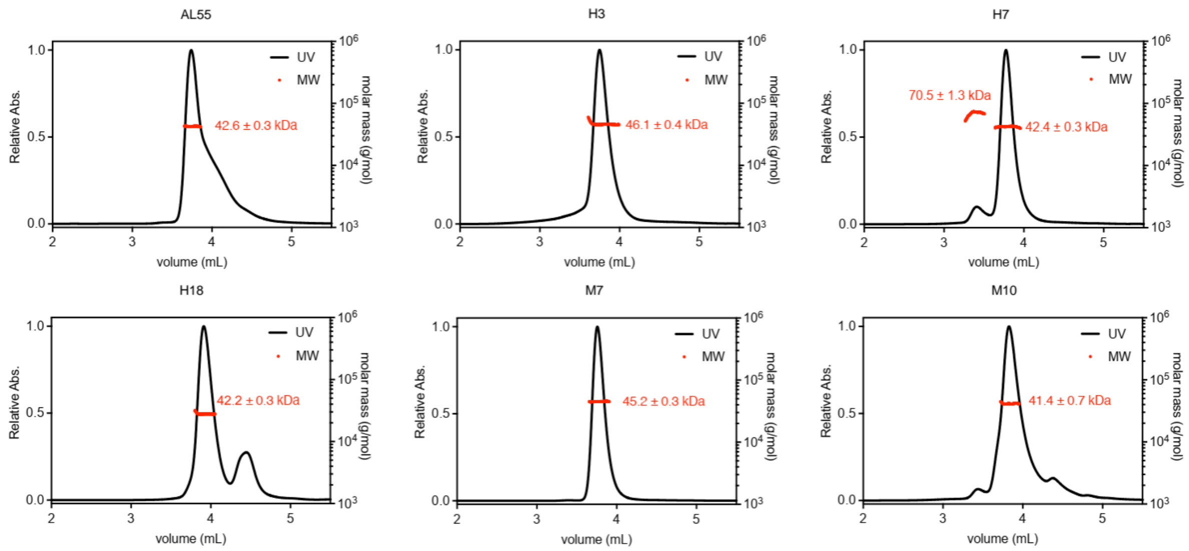


**Figure S20**: SEC-MALS analysis of purified LCs (AL55, H3, H7, H18, M7, and M10). The name of each protein is labelled above their corresponding profile. The obtained molecular weight (MW) is highlighted in the red.

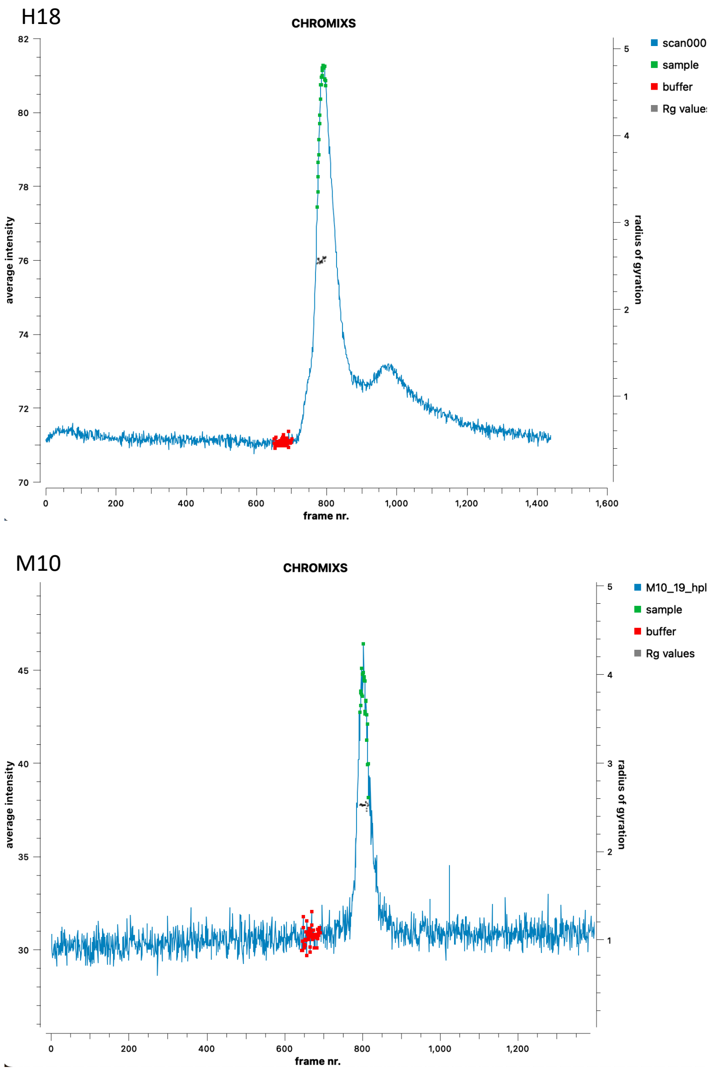


**Figure S21:** SAXS-SEC average intensity for H18 and M10, selected frames for buffer and sample are reported together with the analysis of the estimated Rg values, indicating that the selection corresponds to homogenous conformations.

**Table S1.** Pairwise sequence identity (above diagonal) and similarity (below diagonal) for the 6 systems under study. On the diagonal is reported the germline identified by igBLAST using the IGMT database.

| H3 | H7 | H18 | AL55 | M7 | M10 |
|---|---|---|---|---|---|

| | | | | | | |
|---|---|---|---|---|---|---|
| **H3** | ***IGLV1-44*01*** | 179/216 (82.9%) | 168/216 (77.8%) | 171/218 (78.4%) | 163/216 (75.5%) | 172/217 (79.3%) |
| **H7** | 194/216 (89.8%) | ***IGLV1-51*01*** | 169/214 (79.0%) | 168/218 (77.1%) | 165/214 (77.1%) | 169/217 (77.9%) |
| **H18** | 183/216 (84.7%) | 182/214 (85.0%) | ***IGLV3-19*01*** | 166/218 (76.1%) | **196/214 (91.6%)** | 171/217 (78.8%) |
| **AL55** | 190/218 (87.2%) | 188/218 (86.2%) | 184/218 (84.4%) | ***IGLV6-57*02*** | **164/218 (75.2%)** | 175/218 (80.3%) |
| **M7** | 181/216 (83.8%) | 179/214 (83.6%) | **204/214 (95.3%)** | **182/218 (83.5%)** | ***IGLV3-19*01*** | 169/217 (77.9%) |
| **M10** | 196/217 (90.3%) | 185/217 (85.3%) | 186/217 (85.7%) | 188/218 (86.2%) | 183/217 (84.3%) | ***IGLV2-14*03*** |

**Table S2.** HDX-MS data summary.

| Datasets | H3 | H7 | AL55 | M10 |
|---|---|---|---|---|
| **HDX reaction details** | 1X Phosphate buffer saline in $D_2O$ (pD 7.0), 25°C | | | |
| **HDX time course (min)** | 0, 0.5, 1, 10, 30, 120, and 240 | | | |
| **Back exchange (mean/IQR)** | ND | | | |
| **No. of peptides** | 61 | 50 | 57 | 62 |
| **Sequence coverage (%)** | 98.6 | 92.5 | 98.6 | 99.1 |
| **Average peptide length/ Redundancy** | 13.6/4.16 | 15.8/4.01 | 14.3/4.06 | 15.1/4.69 |
| **Replicates (technical)** | 3 | 3 | 3 | 3 |
| **Repeatability (average SD)** | 0.04 Da | 0.04 Da | 0.04 Da | 0.05 Da |

#### Legends for Movies

**Movie S1 (separate file).** This movie depicts the 3D structure of H3, color-coded by HDX exchnage and rotating 360 degrees.

**Movie S2 (separate file).** This movie depicts the 3D structure of H7, color-coded by HDX exchnage and rotating 360 degrees.

**Movie S3 (separate file).** This movie depicts the 3D structure of AL55, color-coded by HDX exchnage and rotating 360 degrees.

**Movie S4 (separate file).** This movie depicts the 3D structure of M10, color-coded by HDX exchnage and rotating 360 degrees.

#### Legends for Datasets

**Dataset S1 (separate file).**

SAXS data are available on the SASBDB with accession codes:

H3: SASDVL4
H7: SASDVM4
H18: SASDVN4
AL55: SASDVK4
M7: SASDVP4
M10: SASDVQ4

(Preview links)

M10:https://www.sasbdb.org/draft-preview/6174/8nvunm9j0z/
H18:https://www.sasbdb.org/draft-preview/6173/xhgnu1ol62/
M7:https://www.sasbdb.org/draft-preview/6172/2ehc643lbl/
H7:https://www.sasbdb.org/draft-preview/6171/nxc6tcf69p/
H3:https://www.sasbdb.org/draft-preview/6170/fn74eorw3r/
AL55:https://www.sasbdb.org/draft-preview/6156/mryq2shos7/

**Dataset S2 (separate file).**

DOI: 10.5281/zenodo.12731283, https://dx.doi.org/10.5281/zenodo.12731283

Molecular dynamics simulation trajectories and associated statistical weights.
